# Activity-specificity trade-off gives PI5P4Kβ a nucleotide preference to function as a GTP-sensing kinase

**DOI:** 10.1101/2020.06.09.137430

**Authors:** Koh Takeuchi, Yoshiki Ikeda, Miki Senda, Ayaka Harada, Koji Okuwaki, Kaori Fukuzawa, So Nakagawa, Hongyang Yu, Lisa Nagase, Misaki Imai, Mika Sasaki, YuHua Lo, Atsuo T. Sasaki, Toshiya Senda

**Affiliations:** Molecular Profiling Research Center for Drug Discovery, National Institute of Advanced Science and Technology, 2-3-26 Aomi, Koto, Tokyo 135-0063, Japan; Division of Hematology and Oncology, Department of Internal Medicine, University of Cincinnati College of Medicine, Cincinnati, OH 45267, USA; Structural Biology Research Center, Institute of Materials Structure Science, High Energy Accelerator Research Organization (KEK), Tsukuba, Ibaraki 305-0801, Japan; Department of Chemistry and Research Center for Smart Molecules, Rikkyo University, Toshima, Tokyo 171-8501, Japan; School of Pharmacy and Pharmaceutical Sciences, Hoshi University, Shinagawa, Tokyo 142-8501, Japan; Department of Molecular Life Sciences, Tokai University School of Medicine, Isehara 259-1193, Japan; Department of Cancer Biology, University of Cincinnati College of Medicine, OH 45267, USA; Department of Neurosurgery, Brain Tumor Center at UC Gardner Neuroscience Institute, Cincinnati, OH 45267, USA; Institute for Advanced Biosciences, Keio University, Tsuruoka, Yamagata 997-0052, Japan; Department of Accelerator Science, School of High Energy Accelerator Science, SOKENDAI (the Graduate University for Advanced Studies), 1-1 Oho, Tsukuba, Ibaraki 305-0801, Japan; Department of Molecular Genetics, Institute of Biomedical Science, Kansai Medical University, Hirakata, Osaka, 573-1010, Japan; Signal Transduction Laboratory, National Institute of Environmental Health Sciences, National Institutes of Health, Department of Health and Human Services, 111T. W. Alexander Drive, Research Triangle Park, NC 27709, USA

## Abstract

Most kinases function with ATP. However, contrary to the prevailing dogma, phosphatidylinositol 5-phosphate 4-kinase β (PI5P4Kβ) utilizes GTP as a primary phosphate donor with a unique binding mode for GTP. Although PI5P4Kβ is evolved from a primordial ATP-utilizing enzyme, PI4P5K, how PI5P4Kβ evolutionarily acquired the GTP preference to function as a cellular GTP sensor remains unclear. In this study, we show that the short nucleotide base-recognition motif, TRNVF, is responsible for the GTP binding of PI5P4Kβ, and also confers onto PI5P4Kβ an unexpected specificity that extends to inosine triphosphate (ITP) and xanthosine triphosphate (XTP). A mutational study with GTP analogues suggests that the extended specificity is an obligatory consequence to the acquisition of GTP-dependent activity. However, as the cellular concentrations of ITP and XTP are typically negligible, PI5P4Kβ can still function as a GTP sensor, suggesting that the cellular physiological conditions leave room for the functional evolution of PI5P4Kβ.

## INTRODUCTION

Kinases are essential for a variety of cellular processes, including signal transduction, transcription, and metabolism. There is extraordinary diversity in their structure, substrate specificity, and participating pathways. Protein kinases, which represent the largest superfamily consisting of over 500 different distinct genes in the human genome, share a conserved catalytic domain and structural motif that serves for ATP recognition and catalysis (Bossemeyer, 1995; Endicott et al., 2012; Huse and Kuriyan, 2002; Shen et al., 2005; Taylor et al., 1992; Wang and Cole, 2014; Shi et al., 2006). On the other hand, phosphoinositide kinases and inositol phosphate kinases (IP-kinase, including inositol kinases) form distinct families that target the inositol moieties of substrates. Although the families of phosphoinositide and IP-kinases have distinct folds from protein kinases, all these kinases use ATP as the physiological phosphate donor (Clarke and Irvine, 2011; Loijens et al., 1996; Shears and Wang, 2019; Vadas et al., 2011).

The preference for ATP has been experimentally defined for more than 200 kinases (Becher et al., 2013), most of which have a more than 3-fold preference for ATP over GTP based on their affinity values. While GTP is the second-most abundant triphosphorylated nucleotide in cells (0.1-0.5 mM), the affinity difference coupled the higher physiological concentration of ATP (1-5 mM) (Traut, 1994) result in the occupation of kinase catalytic centers by ATP under most cellular physiological conditions. For the recognition, the guanine base cannot interact in the same way as the adenine base in the nucleotide binding pocket, due to the distinct hydrogen donors and acceptors at the 1^st^ and 6^th^ positions of guanine and adenine (Fig. 1A). There are only a few examples, such as casein kinase II (CKII), of kinases that react equally well with GTP and ATP (Niefind et al., 1999). However, the functional relevance of the GTP utilization of these rare kinases remains elusive.

**Figure 1.**
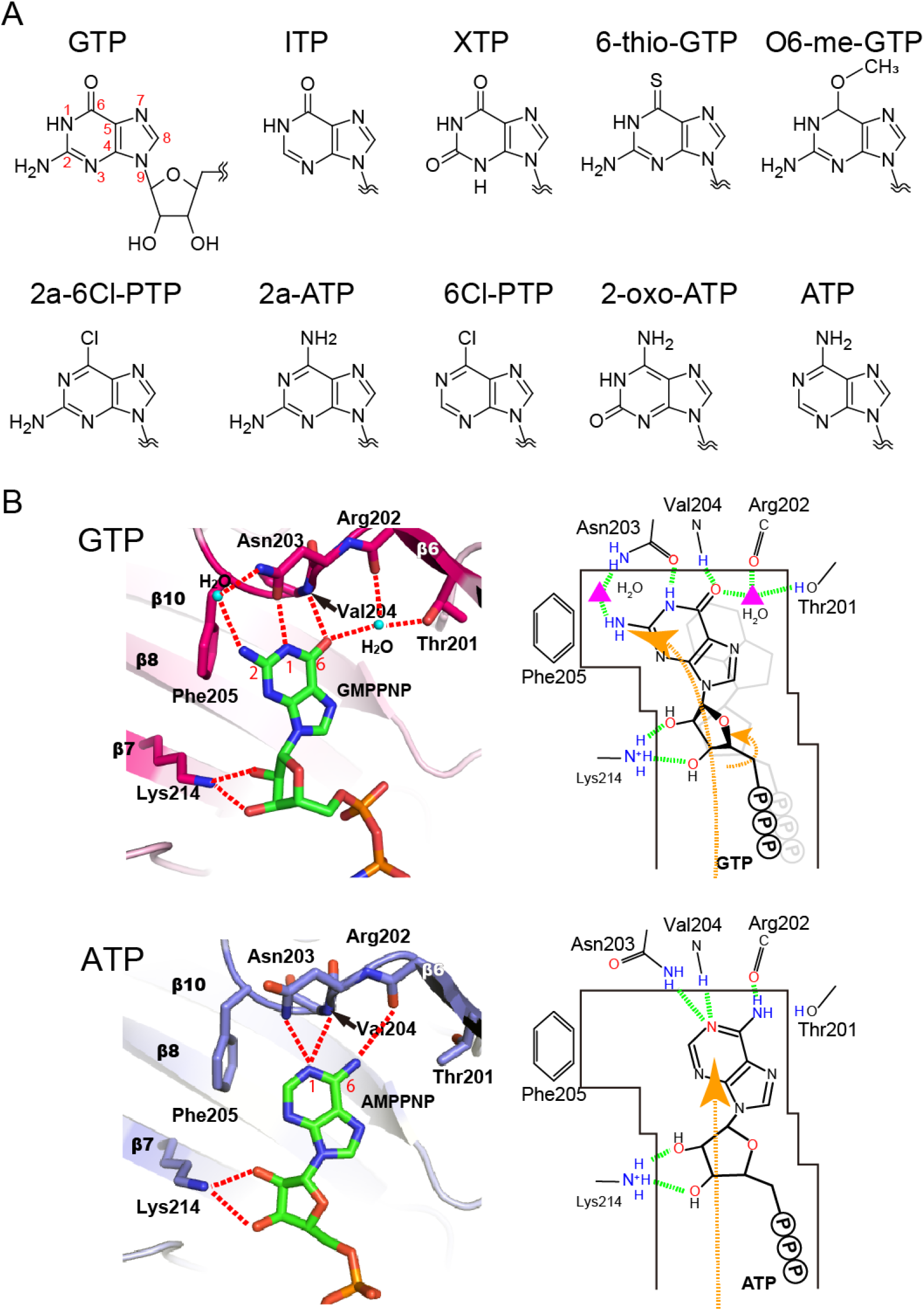
(A) Chemical structures of PNTs used in this study. (B) ATP- and GTP-binding modes of PI5P4Kβ.

The overwhelmingly greater frequency of ATP-preferring kinases has given rise to the dogma that kinases function with ATP. Given this prevailing idea, the strong GTP-preference of phosphatidylinositol 5-phosphate 4-kinase β (PI5P4Kβ) was a surprising discovery (Sumita et al., 2016). PI5P4K, also called Type II PIPK, is a member of the phosphoinositide kinase superfamily and converts the second lipid messenger phosphatidylinositol 5-phosphate (PI(5)P) to phosphatidylinositol 4,5-diphosphate (PI(4,5)P_2_) (Rameh et al., 1997). Despite the higher intracellular concentration of ATP, PI5P4Kβ exhibits a strong preference for GTP and a *K*_M_ value (*K*_M_ for GTP ~88 μM) that is well within the physiological variation of GTP concentration (Sumita et al., 2016). Importantly, a structure-based reverse genetic analysis demonstrated that PI5P4Kβ acts as an intracellular GTP sensor (Sumita et al., 2016; Takeuchi et al., 2016). Interestingly, an evolutionarily cognate phosphoinositide-kinase, PI4P5K/Type I PIPK, utilizes ATP for its reaction (Sumita et al., 2016). A recent report suggests that the divergence of PI5P4K from the PI4P5K family likely occurred at the ancestral lineage of Choanoflagellates and Filasterea (Khadka and Gupta, 2019). The PI5P4K genes are found in a variety of organisms belonging to the Holozoa clade of eukaryotes; however, these genes are not found in the deeper-branching eukaryotic lineages, or in either plants or fungi. Therefore, PI5P4Kβ represents an intriguing example of evolutionary switching of nucleotide preference from ATP to GTP. Considering the high sequence identity between the PI5P4Kβ and PI4P5K subfamilies (>60%), analysis of the amino acid substitutions in the catalytic pocket could uncover the structural requirement that allowed PI5P4Kβ to functionally evolve to an intra-cellular GTP-sensor during the development and homeostasis of multicellular animals.

In this study, we defined the essential interactions that account for GTP- and ATP-binding specificity through an extensive comparison of the GTP- and ATP-binding motifs of kinases and G-proteins in the protein data bank (PDB) (Berman et al., 2003). Then, we biochemically and structurally characterized the nucleotide preference of PI5P4Kβ by a systematic utilization of 10 different purine nucleotide triphosphates (PNTs) (Fig. 1A) and introduction of amino-acid substitutions to the nucleotide-binding pocket. These analyses revealed a trade-off relationship between the GTP dependent activity and nucleotide specificity of PI5P4Kβ. Our results provide a basis critical to understanding how, following a limited and non-detrimental amino acid substitutions in a small stretch of sequences, PI5P4Kβ acquires a GTP preference to function as an intracellular GTP sensor.

## RESULTS

### The ATP recognition mode is shared among protein and lipid kinases

To gain insights into the typical ATP- and GTP-binding modes of proteins and compare them with those of PI5P4Kβ (Fig. 1B), we analyzed 702 unique nucleotide-bound structures for protein kinases, phosphoinositide kinases, inositol phosphate kinases (including inositol kinase), and 134 G-proteins in the PDB. The catalytic domains of the kinases consist of two lobes harboring an ATP-binding site at the hinge region. Binding of an adenine base by kinases is characterized by conservation of two mainchain hydrogen bonds to N(1) and N(6) (Fig. 2A), while other interactions that are unique in each kinase are also observed. Typically, two hydrogen bonds are formed between the mainchain amide and carbonyl groups from the *i*+*2*^*th*^ and *i*^*th*^ residues, respectively. The mode of interaction can be achieved by an extended conformation of the polypeptide and is also conserved in the ATP-binding mode of PI5P4Kβ (Fig. 1B, bottom). In PI5P4Kβ, the N(1) and N(6) of ATP are recognized by the amide group of Val-204 and carbonyl oxygen of Arg-202, respectively. The N(1) of ATP also forms a hydrogen bond with the Nδ of Asn-203. The cognate ATP kinase PI4P5Kα also interacts with ATP in the same binding mode (Fig. 2A, middle) (Muftuoglu et al., 2016).

**Figure 2.**
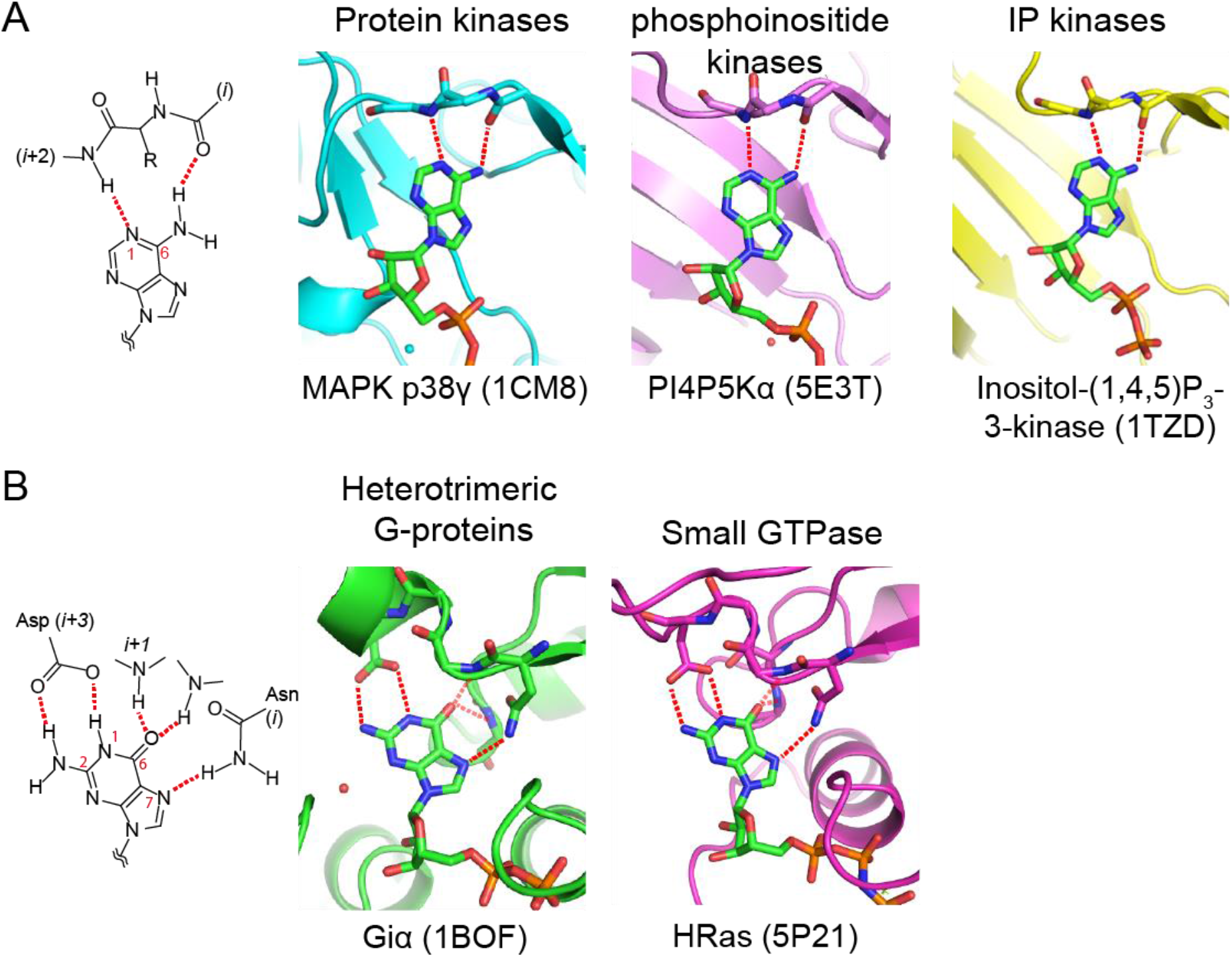
Nucleotide-base binding by kinases and G-proteins. Typical hydrogen-bond interactions between nucleotide bases and proteins are shown for (A) an adenine base in kinases and (B) a guanine base in G-proteins. Representative nucleotide-complex structures from major protein families are also shown.

### The unique GTP-binding mode of PI5P4Kβ by the TRNVF motif

Because the arrangement of a hydrogen donor and acceptor in the guanine base differs from that of the adenine, PI5P4Kβ has a specific GTP-binding mode (Fig. 1B, top). The mode of interaction is different from that of G-proteins, which utilize the conserved NKXD motif for guanine base recognition (Fig. 2B). In G-proteins, the N(1) and NH_2_(2) of the guanine base are simultaneously recognized by the sidechain carboxylate of Asp in the NKX**<u>D</u>** motif (Wennerberg et al., 2005). In addition, in most cases, the N(7) of the guanine base forms a hydrogen bond with Oδ of Asn in the <u>**N**</u>KXD motif, and O(6) forms a hydrogen bond(s) with a neighboring *i*+*1*^*th*^ Lys and a remote mainchain amide group(s).

PI5P4Kβ also forms hydrogen bonds to N(1), NH_2_(2), and O(6) of the guanine ring; however, the interacting residues are distinct from those of the G-protein. PI5P4Kβ utilizes the TRNVF motif (residues 201-205 in humans) to recognize GTP (Fig. S1). Asn-203 in the TR<u>**N**</u>VF motif is structurally located at the corresponding position of the conserved Asp residue of the G-protein, as its Oδ and Nδ atoms form direct and indirect hydrogen bonds with N(1) and NH_2_(2), respectively (Fig. 1B, top). While G-proteins typically have pico to sub-nano molar affinity to GTP, the affinity of PI5P4Kβ to GTP seems to be much weaker, as its *K*_M_ value is only ~100 μM. The indirect hydrogen bond between the Asn-203 Nδ and NH_2_(2), which is mediated by a water molecule, might account for the weaker affinity of PI5P4Kβ compared to that of the G-protein. Another characteristic feature of PI5P4Kβ is a hydrogen-bond network around O(6) of the guanine base involving Thr-201, Arg-202, and Val-204 in the <u>**TR**</u>N<u>**V**</u>F motif and a water molecule (Fig. 1B). These interactions for GTP are enabled by a 1.5 Å shift of the base moiety relative to the ATP. The contribution of the hydrogen-bond network around O(6) for guanine base recognition is also evident from the fragment molecular orbital (FMO) calculation (Fedorov and Kitaura, 2009; Fedorov et al., 2012; Tanaka et al., 2014). Both Val-204 and the water molecule held by Thr-201 show an energetically favored interaction to O(6) of guanine base (Figs. S2A and C). Note that the interaction seems to be even stronger than the aforementioned Asn-203 plus water interactions with the NH_2_(2) position (Figs. S2A and B). The shift of the base position promotes a formation of aromatic-aromatic interactions with Phe-205 in the TRNV<u>**F**</u> motif (Fig. S2A), which is unique to guanine base recognition. Interestingly, the guanine and adenine base recognition of CKII and PI5P4Kβ has similarity in the hydrogen-bond networks around N(1) and O(6) as well as the 1.5 Å shift of guanine base compared to that of the adenine ring (Figs. 1B and S3) (Niefind et al., 1999); however, CKII uses only mainchain atoms for base recognitions.

The GTP-recognizing TRNVF sequence also serves for the adenine-base recognition and is strictly conserved among PI5P4Kβ proteins (Fig. S1). Therefore, the TRNVF sequence can be designated as a dual nucleotide base-binding motif. Especially, Thr-201, Asn-203, and Phe-205 in the motif would be of importance as their sidechains contribute to the interaction with the guanine base. In contrast, among the ancestral ATP-dependent PI4P5Ks, the MNNψL sequence is conserved, where “ψ” is donated for branched amino acids (Fig. S1). This indicates that PI5P4Kβ has established an atypical mode of GTP recognition, while conserving the canonical ATP-binding mode, by changing a few residues in the MNNψL sequence into the TRNVF motif. Especially, Thr-201 and Phe-205, which establish sidechain interactions with the guanine base, would be of importance due to their unique contributions to the guanine-base recognition.

### PI5P4Kβ can hydrolyze XTP and ITP

Next, the mechanism of the GTP preference of PI5P4Kβ was investigated using a series of ATP and GTP analogs. Based on analysis of the GTP- PI5P4Kβ interaction (Fig. 1B), 10 PNTs with different configurations at the 2^nd^ and 6^th^ positions of the purine base (NH_2_(2) and O(6) in guanine base, respectively) were chosen (Fig. 1A). The hydrolysis activities of PI5P4Kβ for these PNTs were quantified by an NMR-based assay (Fig. S4). The intrinsic hydrolysis activity of PI5P4Kβ (*i.e.,* the transfer of phosphoryl to water, instead of PI(5)P) has been shown to reflect the characteristic GTP-preference of the kinase (Sumita et al., 2016). PI5P4Kβ showed substantial activity with ITP, XTP, 6-Thio-GTP, and 2a-ATP (Fig. 3A), indicating that NH_2_(2) is dispensable for the activity of GTP-like PNTs, since both ITP and XTP lack the NH_2_(2) moiety. On the other hand, O(6) seems to be required for the activity. ITP and XTP, both of which have the O(6) moiety, showed 1.3- and 1.9-times higher hydrolysis activity compared to GTP, but O6-me-GTP and 2a-6Cl-PTP, which lack the O(6) moiety, showed very low hydrolysis activity. In line with this notion, 6-thio-GTP, which possesses sulfate, which is structurally and electrostatically similar to oxygen in the 6th position, can also be utilized by PI5P4Kβ (Fig. 3A). A competition assay between these PNTs and GTP showed that the GTP-dependent PI(5)P phosphorylation activity was strongly inhibited by ITP, XTP, and 6-thio-GTP (Fig. S5), supporting the idea that the specificity of PI5P4Kβ extends beyond GTP due to the strong dependence on the O(6) interaction in the nucleotide recognition. This view is also supported by the larger energetic contribution of the O(6) moiety compared to the NH_2_(2) moiety in the FMO interaction analysis between GTP andPI5P4Kβ (Fig. S2).

**Figure 3.**
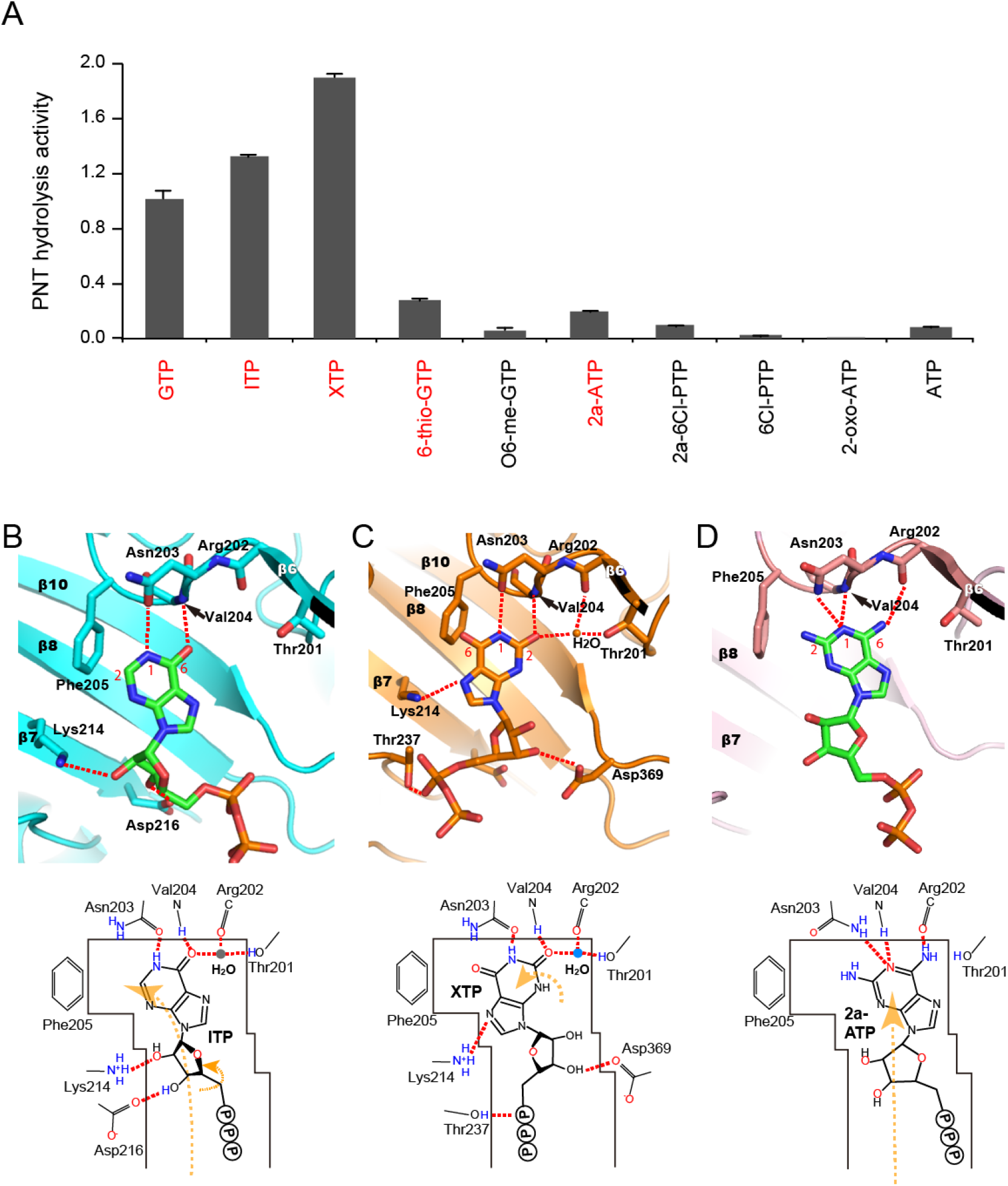
PNT hydrolysis activity and binding modes of PI5P4Kβ. (A) PNT hydrolysis activity of PI5P4Kβ. The PNT hydrolysis activity of PI5P4Kβ was assessed by the signal intensity ratios of diphosphorylated/triphosphorylated nucleotides after the reaction. The average values from three experiments are shown with error bars (S.D.). Highly hydrolyzed nucleotides (> 0.1 of the ratios) are indicated by red. (B, C, and D) Interactions of the PI5P4Kβ-ITP, PI5P4Kβ-XTP, and PI5P4Kβ-2a-ATP complexes, respectively. Red dotted lines represent hydrogen bonds between the PNTs and PI5P4Kβ. In the case of XTP, only the XTP-binding mode is shown. As for the ITP complex, the position of the water that is visible with a lower σ value is also indicated with a gray circle.

### Crystal structures of PI5P4Kβ unveil the recognition mechanism of the active triphosphorylated nucleotides

Next, we analyzed mechanistic details for the extended specificity of PI5P4Kβ beyond GTP using the crystal structures of PI5P4Kβ complexed with any of three PNTs: ITP, XTP, or 2a-ATP (Figs. 3B-D, S6-7, and Table S1). Since the soaking of the 6-thio-GTP broke PI5P4Kβ crystals, the crystal structure of the 6-this-GTP complex could not be obtained. In the 2a-ATP complex, 2a-ATP binds to PI5P4Kβ with a binding mode similar to that of ATP (Fig. 3D), except that the N(2) of 2a-ATP and the Phe-205 sidechain seem to form an additional van der Waals interaction. This additional interaction would explain the stronger binding of 2a-ATP compared to ATP. The crystal structure of the ITP complex revealed that the interaction with the inosine base is essentially the same as that with the guanine base (Figs. 3B and S6A). The water molecule that participates in the hydrogen-bond network around O(6) is less clear in the PI5P4Kβ-ITP complex; however, the presence of the water molecule is evident when the criterion for identifying it is slightly lowered (2σ). Since ITP, which lacks NH_2_(2), can reside in the G-site, the contribution of NH_2_(2) to the GTP binding to PI5P4Kβ would be minor. Nevertheless, the absence of an interaction with NH_2_(2) slightly changes the position of the nucleotide base of ITP relative to GTP, which might explain why the hydrolysis activity of ITP is higher than that of GTP, since the position of the nucleotide base affects the phosphate group positions in the catalytic site.

Surprisingly, XTP has two different but overlapping binding modes in the binding site of PI5P4Kβ. In the first binding mode, XTP is in the G-site forming hydrogen bonds of N(1) and O(6) corresponding to those found in GTP (Fig. S8). An indirect hydrogen bond between O(2) and Asn-203 via water was not observed. In the second binding mode, the base of the XTP is flipped by 180° respective to the first binding mode, revealing the distinct XTP-binding mode (Fig. 3C). Even after the base flip, XTP forms a hydrogen-bond network similar to the first one; the N(1) occupies an almost identical position within 1 Å difference, and the positions of O(2) and O(6) are merely swapped. As a result, the N(1) still forms a hydrogen bond with Asn-203 Oδ, as observed in the GTP-binding mode. The O(2) of XTP forms bifurcated hydrogen bonds to the mainchain amide group of Val-204 and a water molecule, which in turn forms a hydrogen bond with Oγ of Thr-201 (Fig. 3C). The presence of these two distinct binding modes for XTP would explain the elevated activity of PI5P4Kβ on XTP. Nevertheless, these structural studies showed that ITP and XTP are GTP-type PNTs, in which the hydrogen-bond network around O(6) is critical for the interaction.

### Rational PI5P4Kβ mutants define the contribution of key residues to the GTP, ATP, and XTP-binding

To analyze the contribution of the nucleotide interacting residues to the GTP-, ATP-, and XTP-binding modes, we compared the effect of mutations of Thr-201, Asn-203, and Phe-205 in the TRNVF motif (Fig. 4). Since Thr-201 and Phe-205 are substituted to Met and Leu in PI4P5K (or Type I PIPK) (Fig. S1), respectively, the T201M and F205L mutants could provide insight into the evolutionary change of the base-specificity of PI5P4K.

**Figure 4.**
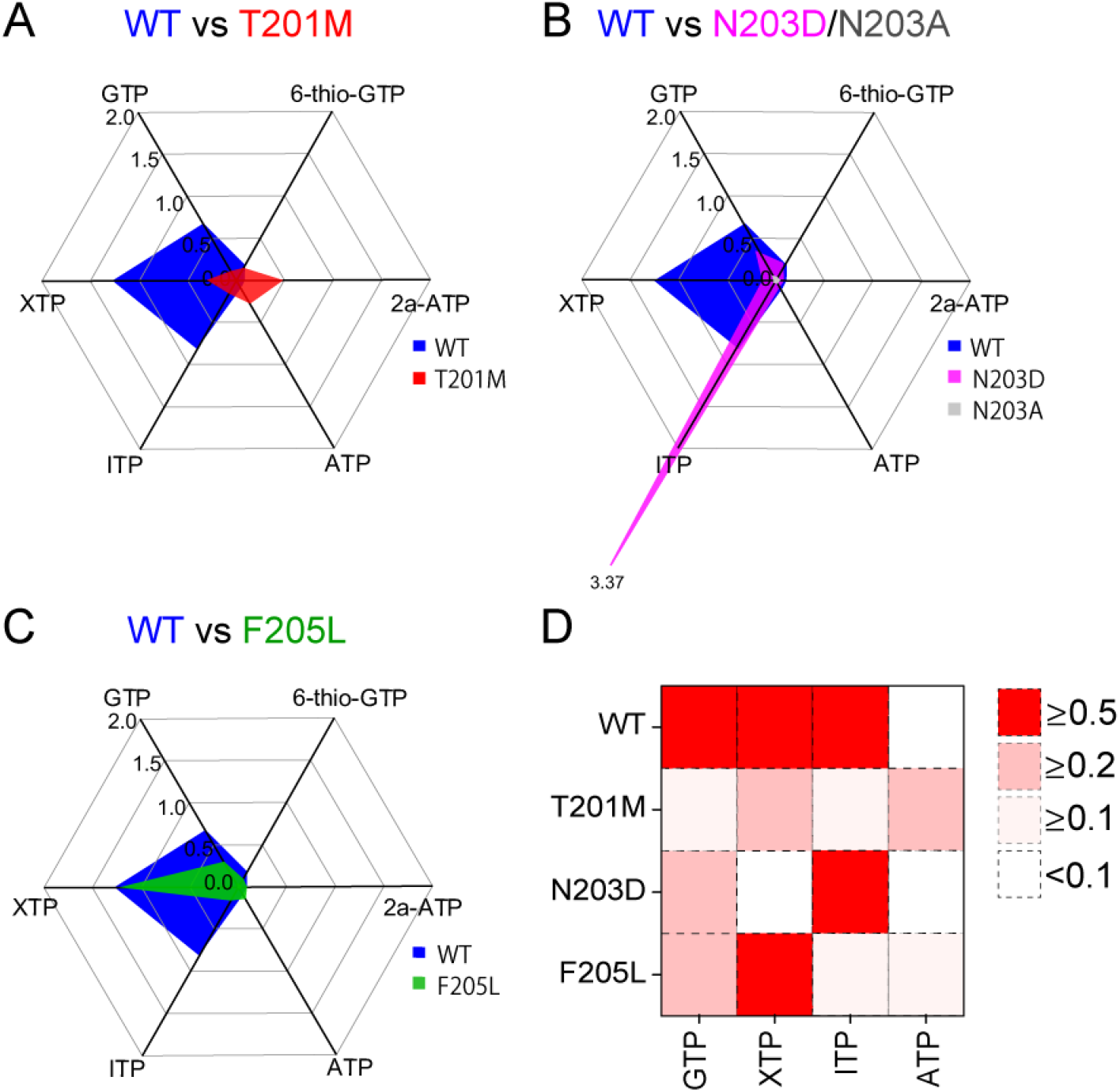
GTP-, ATP-, and XTP-binding modes and effect of mutations on PI5P4Kβ activity for different PNTs. **(A-C)** PNTs hydrolysis activity of mutant PI5P4Kβ as compared to the WT. The ratios of dephosphorylated/triphosphorylated nucleotides after reaction are shown. The N203A activity is almost negligible, and thus, it is barely visible in the figure. (D) The ratios of dephosphorylated/triphosphorylated nucleotides after reaction are shown in color depth to compare PNTs hydrolysis activity and specificity among the WT and mutant PI5P4Kβs.

We first analyzed the functional role of Asn-203, because this is the only invariant residue in the base-recognition loop of PI4P5K and PI5P4K. The mutation of this residue markedly reduced the binding to both ATP and GTP (Fig. 4B; gray). The FMO calculation clearly showed the importance of this interaction; approximately 1/4 of the interaction energy between the nucleotide base and PI5P4Kβ is contributed by a hydrogen bond between the guanine base and Asn-203 (Fig. S2).

The effects of the T201M (PI5P4Kβ^T201M^) and F205L (PI5P4Kβ^F205L^) mutations have already been partly reported (Sumita et al., 2016). The decreased GTP-dependent kinase activity of PI5P4Kβ^T201M^ has been explained by the loss of the hydrogen-bond network around O(6) by the mutation (Fig. 1B), and the higher ATP-dependent activity of PI5P4Kβ^T201M^ seems to have arisen from an additional hydrophobic interaction between the adenine-base and the substituted Met (Sumita et al., 2016). In the case of PI5P4Kβ^F205L^, the loss of the π-π interaction between the guanine base and the aromatic ring of Phe-205 caused the reduction of the GTP-dependent activity. In contrast, the ATP-binding was not affected by the F205L mutation, and PI5P4Kβ^F205L^ retains an ATP-dependent activity comparable to that of WT PI5P4Kβ (Sumita et al., 2016).

In addition, we analyzed the hydrolysis activity of the mutants on four *active* triphosphorylated nucleotides, XTP, ITP, 6-thio-GTP, and 2a-ATP. As expected, PI5P4Kβ^T201M^ was substantially less active on the GTP-type PNTs (Figs. 4A and D), showing the importance of the hydrogen-bond network around O(6). In contrast, PI5P4Kβ^T201M^ was more active on 2a-ATP and ATP, which shares the ATP-mode interaction (Figs. 1A and S9A), as these nucleotide bases show additional interactions with the mutated methionine sidechain. Intriguingly, PI5P4Kβ^N203D^ and PI5P4Kβ^F205L^ showed much stronger hydrolysis activity for a single NTP other than GTP (Figs. 4B and C). PI5P4Kβ^N203D^ is hyperactive to ITP, while the activities on GTP, XTP, ATP, and 2a-ATP were significantly reduced (Fig. 4B and D). In the crystal structure of the PI5P4Kβ^N203D^-ITP complex, ITP binds in a similar manner as the WT (Figs. 3B and S9B). Although the crystal structure could not explain the hyper ITPase activity of PI5P4Kβ^N203D^, the results suggest the importance of recognizing the 1^st^ position in both GTP- and ATP-mode interactions.

The Phe-205 to Leu mutation makes a protein less active on GTP, ITP, and 6-thio-GTP (Figs. 4C and D), suggesting the importance of the π-π interaction between the Phe-205 sidechain and the nucleotide bases in the GTP-binding mode (Fig. S2). The diminished susceptibility of XTP might be due to the presence of additional binding modes (the XTP-binding mode), which make the nucleotide less susceptive to the F205L mutation (Fig. S9C). It should also be noted that the activity on ATP and 2A-ATP were not affected by the F205L mutation, as the sidechain did not contribute to the interaction in the ATP-binding mode (Sumita et al., 2016).

## Discussion

Our analyses provide a novel insight into the molecular evolution by which PI5P4Kβ acquired a GTP preference. Human PI5P4Kβ has a ^201^TRNVF^205^ motif as the dual base-recognition fragment that is conserved among PI5P4Kβs from very ancient vertebrates, such as elephant sharks and coelacanths, to humans (Fig. S1) (Sumita et al., 2016). While all residues in the base-recognition fragment are involved in the specific interaction with the guanine base in PI5P4Kβ (Fig. 1B), PI5P4Kβ has evolved functional roles for the sidechains of the 1^st^ and 5^th^ residues in the base-recognition <u>**T**</u>RNV<u>**F**</u> motif to achieve the interaction with the guanine base necessary to function as a GTP sensor. Our biochemical analysis revealed an unexpected specificity of WT PI5P4Kβ extending to ITP and XTP (Fig. 3); these NTPs structurally share O(6) as a proton acceptor. The FMO analysis revealed that the hydrogen-bonding interaction with the O(6) position by Val-204 and a water molecule stabilized by Thr-201 play critical roles in the base recognition (Fig. S2). Indeed, PI5P4Kβ^T201M^ shows a substantially reduced hydrolysis activity for GTP and its analogues. The O(6) position of the nucleotide bases is the site that chemically differentiates ATP from GTP; thus the strong interaction with PI5P4Kβ is reasonable for its acquisition of the GTP preference. However, the benefit only comes with the extended specificity to XTP and ITP. Phe-205 also contributes to the extended specificity via π-π interaction. These results suggest that there is a trade-off in the relationship between the GTP-dependent activity and nucleotide specificity in the WT PI5P4Kβ. The kinase seems to gain the benefit to function with GTP without much sacrifice, as the intracellular concentrations of ITP and XTP are far smaller than that of GTP *in vivo*. Thus, the parasitic activity on XTP and ITP is not detrimental for the biological function of PI5P4Kβ.

In the course of evolution, PI5P4Kβ made a compromise to acquire the cellular GTP-sensing function under the given environment and the restraint of the phosphoinositide kinase fold. Considering that the TRNVF motif of the PI5P4Kβ isotype is conserved among vertebrates (Fig. S1), this appears to be the strongly favored evolutional solution for the GTP recognition, within the premised structure and sequence of the phosphoinositide kinases. It should be noted that neither PI5P4Kβ^T201M^ nor PI5P4Kβ^F205L^ loses its original ATP-dependent activity (Fig. 4). Therefore, having a *semi-optimal* sequence between the TRNVF motif and the original MNNψL motif might not be detrimental, as we successfully established a cell line that only has PI5P4Kβ^F205L^ in our previous paper (Sumita et al., 2016). Such non-detrimental mutations might allow a smooth evolutional transition of an ATP-dependent kinase to a GTP-sensing kinase. Given the importance of the GTP concentration in controlling energy metabolism and proliferation (Kofuji et al., 2019), the establishment of a new mode of signaling that utilizes GTP through the evolution of PI5P4Kβ or, more specifically, through two critical amino acid substitutions in the TRNVF motif, would be of critical importance for energy homeostasis in higher animals and would represent a potential mechanism for gaining additional functionality during evolution.

## Acknowledgement

We thank Ms. Tomomi Sato for her contribution in the early stage of this work. We thank Ms. Emily Dobbs, Dr. Yuki Fujii, and Dr. Eric P. Smith for excellent proofreading.

## Material and Methods

### Materials

All materials were purchased from Sigma-Aldrich or Fujifilm Wako Pure Chemical Corporation, unless otherwise noted. Phospholipids were purchased from Echelon Biosciences. PNTs were purchased from TriLink BioTechnologies

### Buffers

Buffer A: 50 mM Tris-HCl (pH 8.0), 300 mM NaCl, 5 mM β-mercaptoethanol (βMe), and 10 mM imidazole
Buffer B: 50 mM Tris-HCl (pH 7.5), 300 mM NaCl, 5 mM βMe, and 300 mM imidazole
Buffer C: 50 mM Tris-HCl (pH 8.0), 100 mM NaCl, and 2 mM DTT
Crystallization buffer (reservoir solution): 100 mM sodium-citrate (pH5.5), 10 mM magnesium-acetate, 100 mM lithium-acetate, and 10% (v/v) polyethyleneglycol-4000 (PEG-4000)
NMR buffer: 10 mM sodium phosphate (pH 6.8), 100 mM NaCl, 10 mM MgCl_2_, 2 mM DTT, and 99.6% D_2_O

### Expression vectors

Expression vectors of human PI5P4Kβs were generated by the standard polymerase chain reaction (PCR) method as described by Sumita et al. (2016). The full-length PI5P4Kβ was used in biochemical and NMR assays, while the first 31 amino-acid residues of PI5P4K were further deleted by QuikChange mutagenesis for X-ray crystallography as described by Sumita et al. (2016). The N-terminal truncated PI5P4Kβ (a.a. 32-416) showed the same enzymatic activity as the full-length PI5P4Kβ; however, it gave crystals with better diffraction. The T201M-, N203D-, N203A-, and F205L-PI5P4Kβ mutants were generated using primer pairs 1, 2, 3, and 4, respectively.

### Primers

Primer pair 1: T201M 5’-ATGGTGGTTATGAGGAACGTGTTC-3’ 5’-CACGTTCCTCATAACCACCATGTA-3’
Primer pair 2: N203D 5’-GGTGGTTACCAGGGACGTGTTCAGCCATC-3’ 5’-GATGGCTGAACACGTCCCTGGTAACCACC-3’
Primer pair 3: N230A 5’-CATGGTGGTTACCAGGGCGGTGTTCAGCCATCGG-3’ 5’-CCGATGGCTGAACACCGCCCTGGTAACCACCATG-3’
Primer pair 4: F205L 5’-CCAGGAACGTGTTGAGCCATCGG-3’ 5’-ATGGCTCAACACGTTCCTGGTAACC-3’

### Expression and purification of PI5P4Kβ

The human PI5P4Kβ was expressed and purified as described previously (Sumita et al., 2016). Briefly, the BL21 (DE3) harboring the PI5P4Kβ expression vector was inoculated in 5 ml of LB medium containing 50 μg/ml of kanamycin at 37 °C overnight. Cells were collected by centrifugation and further inoculated into 1 liter of Luria-Bertani medium. When the bacterial culture was grown to OD_600_ = ~0.6, 0.6 mM of isopropyl-β-D-thiogalactopyranoside was added to induce the protein overexpression. The induced culture was further incubated at 25 °C for 16 h and harvested with centrifugation.

The cells were lysed by sonication in the Buffer A. The lysate was centrifuged to remove cell debris and the PI5P4Kβ protein was purified from the supernatant of the cell lysate by Ni-affinity chromatography. The cell lysate was applied on 2 ml of Ni-NTA agarose (QIAGEN) equilibrated with Buffer A. After applying the cell lysate, the column was washed with Buffer A and the PI5P4Kβ protein was eluted from the column with buffer B. The elution was concentrated to about 10 ml by ultrafiltration using Amicon Ultra with MWCO of 10K (Millipore), and 80 μl of PreScission protease was added per 10 mg of PI5P4Kβ. The solution was dialyzed against Buffer C overnight at 4 °C and applied on a Resource Q column (GE Healthcare) that was equilibrated with Buffer C. PI5P4Kβ was eluted with a linear gradient of 30 column volumes and an increasing ionic strength up to 400 mM NaCl. The fractions corresponding to the PI5P4Kβ dimer, which would be eluted around 250 mM NaCl concentration, were collected and concentrated to 20 mg/ml and stored at −80 °C. The expression and purification procedures of PI5P4Kβ mutants were essentially identical to those for the WT PI5P4Kβ.

### NMR spectroscopy

All experiments were performed on Bruker Avance 700 MHz spectrometer equipped with a triple resonance probe. All spectra were collected in the NMR buffer (see buffer list above) at 25 °C. Spectra were processed using TOPSPIN (Bruker).

For the PNT-hydrolysis assay, 250 μM PNTs were mixed with 2 μM PI5P4Kβ and hydrolysis reactions were carried out at 25 °C for 20 h. The reaction was monitored by the ratio of the intensity of the H8 position signal from the dinucleotide- over trinucleotide-forms in the 1D ^1^H experiments.

### Crystallization, data collection, and structure determination

PI5P4Kβ was crystallized with the sitting-drop vapor diffusion method by mixing 1.5 μl of protein solutions (20 mg/ml) and 1.5 μl of crystallization buffer (reservoir solution) (see buffers in the supplemental materials) at 4°C; these conditions were optimized based on the previously reported crystallization conditions (Sumita et al., 2016). Under the optimized conditions, plate-shape crystals with approximate dimensions of 0.4 × 0.4 × 0.1 mm^3^ appeared in 1 - 2 days.

To prepare PI5P4Kβ-PNT complex crystals, we applied the double-soaking method (Senda et al., 2016). In the first step, apo PI5P4Kβ crystals were soaked in a reservoir solution containing 15% (w/v) PVP-K15 and 1 mM of PNTs for 20 h at 20 °C. The crystals were then transferred to another reservoir solution supplemented with 12.5% (w/v) PVP-K15, 15% (v/v) ethyleneglycol (EG), and 1 mM of the PNTs for 30 sec.

Diffraction data were collected at beamline BL17A and NE3A of the PF in KEK (Tsukuba, Japan). The wavelength and temperature were set to 0.9800 Å or 1.0000 Å and −178 °C, respectively. The diffraction data were processed and scaled using the programs XDS and XSCALE, respectively (Kabsch, 2010). The apo-PI5P4Kβ crystal before soaking belonged to space group *C*222_1_, with unit-cell parameters *a* = 107.1 Å, *b* = 182.4 Å, *c* = 105.4 Å.

The structures of the PI5P4Kβ-PNT complexes were determined by the molecular replacement method using the PHENIX program (Liebschner et al., 2019). The structure of human PI5P4Kβ (PDB ID: 3WZZ) in the *apo* form was used as a search model (Sumita et al., 2016). A simulated annealing m*Fo*-D*Fc* omit map corresponding to the individual nucleotide was used to confirm the conformation of the bound nucleotides (Fig. S8). Water molecules were modeled only for those nucleotides clearly visible (> 3σ) in m*Fo*-D*Fc* maps. Crystallographic refinement was performed by the phenix.refine program (Liebschner et al., 2019).

Data collection and structure refinement statistics are listed in Table S1. For WT-PI5P4Kβ, the favored Ramachandran values were, 89.1%, 91.7%, 92.6%, 90.0%, and 89.9% for the apo, WT-GMPPNP, WT-AMPPNP, WT-ITP, WT-XTP, and WT-2A-a-ATP complexes, respectively. For the PI5P4Kβ mutants, the favored Ramachandran values were, 91.1%, 90.7%, 91.3%, 91.0%, 88.9%, and 89.7% for the T201M-2a-ATP, N203D-ITP, N203D-XTP, F205L-ITP, and F205L-XTP complexes, respectively. There were no disallowed Ramachandran values. Molecular graphics were prepared by PyMOL (Schrödinger, LLC.).

All atomic coordinates obtained in the present study have been deposited in the PDB under the accession codes as shown in the statistics table.

### Structural analysis of kinases and G-proteins

702 protein kinase, PI-kinase, and IP-kinase (including inositol kinase) structures in complex with any types of ATP and ATP analogs, including ATPγS, AMPPNP, AMPPCP, and ADP, were collected from PDB and the hydrogen-bonding interaction of the kinases that are described in the repository were analyzed. As for G-proteins, 128 small G-proteins and 6 α-subunits of heterotrimeric G-proteins in complex with any types of GTP and GTP analogs, including GTPγS, GMPPNP, GMPPCP, and GDP, were collected from the PDB and the hydrogen-bonding interaction of the G-proteins that are described in the repository were analyzed. The structural data used in this study are listed in the Supplementary Materials.

### Fragment molecular orbital (FMO) calculations

*Ab initio* FMO calculation (Fedorov and Kitaura, 2009; Fedorov et al., 2012; Tanaka et al., 2014) was performed on the crystal structure of the PI5P4Kβ-GTP complex. While PI5P4Kβ is a homodimer, only the monomeric structure was utilized for the FMO calculation and inter-subunit interactions were neglected. The crystal structure was modified before performing the FMO calculation. First, all water molecules except those interacting with the nucleotide base were deleted from the crystal structure. Second, hydrogen atoms were added based on the assignment of the protonation state that was calculated using the Protonate 3D function of the Molecular Operating Environment program package (Chemical Computing Group, Montreal, Canada). Third, the energy of hydrogen atoms was minimized with the Amber10:EHT force field. Then, FMO calculations for the PI5P4Kβ structure were performed using ABINIT-MP software (Mochizuki et al., 2019; Nakano et al., 2006). The second-order Møller-Plesset perturbation (MP2) (Mochizuki et al., 2004a; Mochizuki et al., 2004b) method was used with the 6-31G* basis function as a theoretical calculation level; namely, the FMO-MP2/6-31G* level of theory was used. For the FMO calculation, PI5P4Kβ was fragmented into amino acid units at bonds between the C and Cα atoms of the mainchain. GTP was fragmented into the base, sugar, and phosphate groups. Each water molecule was treated as a single fragment. This fragmentation treatment made it possible to easily calculate the electronic structure of the whole complex and the IFIEs.

## References

Becher, I., Savitski, M.M., Savitski, M.F., Hopf, C., Bantscheff, M., and Drewes, G. (2013). Affinity Profiling of the Cellular Kinome for the Nucleotide Cofactors ATP, ADP, and GTP. ACS Chemical Biology 8, 599–607.

Berman, H., Henrick, K., and Nakamura, H. (2003). Announcing the worldwide Protein Data Bank. Nature Structural & Molecular Biology 10, 980–980.

Bossemeyer, D. (1995). Protein kinases--structure and function. FEBS Lett 369, 57–61.

Clarke, J.H., and Irvine, R.F. (2011). The activity, evolution and association of phosphatidylinositol 5-phosphate 4-kinases. Adv Enzyme Regul.

Doughman, R.L., Firestone, A.J., and Anderson, R.A. (2003). Phosphatidylinositol phosphate kinases put PI4,5P(2) in its place. J Membr Biol 194, 77–89.

Endicott, J.A., Noble, M.E.M., and Johnson, L.N. (2012). The Structural Basis for Control of Eukaryotic Protein Kinases. Annual Review of Biochemistry 81, 587–613.

Fedorov, D., and Kitaura, K. (2009). The Fragment Molecular Orbital Method: Practical Applications to Large Molecular Systems (CRC Press).

Fedorov, D.G., Nagata, T., and Kitaura, K. (2012). Exploring chemistry with the fragment molecular orbital method. Physical Chemistry Chemical Physics 14, 7562–7577.

Huse, M., and Kuriyan, J. (2002). The conformational plasticity of protein kinases. Cell 109, 275–282.

Kantardjieff, K.A., and Rupp, B. (2003). Matthews coefficient probabilities: Improved estimates for unit cell contents of proteins, DNA, and protein-nucleic acid complex crystals. Protein science: a publication of the Protein Society 12, 1865–1871.

Khadka, B., and Gupta, R.S. (2019). Novel Molecular Signatures in the PIP4K/PIP5K Family of Proteins Specific for Different Isozymes and Subfamilies Provide Important Insights into the Evolutionary Divergence of this Protein Family. Genes (Basel) 10.

Kofuji, S., Hirayama, A., Eberhardt, A.O., Kawaguchi, R., Sugiura, Y., Sampetrean, O., Ikeda, Y., Warren, M., Sakamoto, N., Kitahara, S., et al. (2019). IMP dehydrogenase-2 drives aberrant nucleolar activity and promotes tumorigenesis in glioblastoma. Nat Cell Biol 21, 1003–1014.

Liebschner, D., Afonine, P.V., Baker, M.L., Bunkoczi, G., Chen, V.B., Croll, T.I., Hintze, B., Hung, L.-W., Jain, S., McCoy, A.J., et al. (2019). Macromolecular structure determination using X-rays, neutrons and electrons: recent developments in Phenix. Acta Crystallographica Section D 75, 861–877.

Loijens, J.C., Boronenkov, I.V., Parker, G.J., and Anderson, R.A. (1996). The phosphatidylinositol 4-phosphate 5-kinase family. Adv Enzyme Regul 36, 115–140.

Matthews, B.W. (1968). Solvent content of protein crystals. J Mol Biol 33, 491–497.

Mochizuki, Y., Koikegami, S., Nakano, T., Amari, S., and Kitaura, K. (2004a). Large scale MP2 calculations with fragment molecular orbital scheme. Chemical Physics Letters 396, 473–479.

Mochizuki, Y., Nakano, T., Koikegami, S., Tanimori, S., Abe, Y., Nagashima, U., and Kitaura, K. (2004b). A parallelized integral-direct second-order Møller–Plesset perturbation theory method with a fragment molecular orbital scheme. Theoretical Chemistry Accounts 112, 442–452.

Mochizuki, Y., Nakano, T., Okiyama, Y., Sakakura, K., Akinaga, Y., Watanabe, H., Kato, K., Sato, S., Yamamoto, J., Yamashita, Y., et al. (2019). ABINIT-MP - Open Ver.1. In University of California, San Francisco.

Muftuoglu, Y., Xue, Y., Gao, X., Wu, D., and Ha, Y. (2016). Mechanism of substrate specificity of phosphatidylinositol phosphate kinases. Proceedings of the National Academy of Sciences 113, 8711.

Nakano, T., Mochizuki, Y., Fukuzawa, K., Amari, S., and Tanaka, S. (2006). CHAPTER 2 - Developments and applications of ABINIT-MP software based on the fragment molecular orbital method. In Modern Methods for Theoretical Physical Chemistry of Biopolymers, E.B. Starikov, J.P. Lewis, and S. Tanaka, eds. (Amsterdam: Elsevier Science), pp. 39–52.

Niefind, K., Pütter, M., Guerra, B., Issinger, O.G., and Schomburg, D. (1999). GTP plus water mimic ATP in the active site of protein kinase CK2. Nat Struct Biol 6, 1100–1103.

Rameh, L.E., Tolias, K.F., Duckworth, B.C., and Cantley, L.C. (1997). A new pathway for synthesis of phosphatidylinositol-4,5-bisphosphate. Nature 390, 192–196.

Senda, M., Hayashi, T., Hatakeyama, M., Takeuchi, K., Sasaki, A.T., and Senda, T. (2016). Use of Multiple Cryoprotectants to Improve Diffraction Quality from Protein Crystals. Crystal Growth & Design 16, 1565–1571.

Shears, S.B. (2004). How versatile are inositol phosphate kinases? Biochem J377, 265–280.

Shears, S.B., and Wang, H. (2019). Inositol phosphate kinases: Expanding the biological significance of the universal core of the protein kinase fold. Adv Biol Regul 71, 118–127.

Shen, K., Hines, A.C., Schwarzer, D., Pickin, K.A., and Cole, P.A. (2005). Protein kinase structure and function analysis with chemical tools. Biochim Biophys Acta 1754, 65–78.

Shi, Z., Resing, K.A., and Ahn, N.G. (2006). Networks for the allosteric control of protein kinases. Current Opinion in Structural Biology 16, 686–692.

Sumita, K., Lo, Y.-H., Takeuchi, K., Senda, M., Kofuji, S., Ikeda, Y., Terakawa, J., Sasaki, M., Yoshino, H., Majd, N., et al. (2016). The Lipid Kinase PI5P4K β; Is an Intracellular GTP Sensor for Metabolism and Tumorigenesis. Molecular Cell 61, 187–198.

Takeuchi, K., Senda, M., Lo, Y.H., Kofuji, S., Ikeda, Y., Sasaki, A.T., and Senda, T. (2016). Structural reverse genetics study of the PI5P4Kβ-nucleotide complexes reveals the presence of the GTP bioenergetic system in mammalian cells. FEBS J 283, 3556–3562.

Tanaka, S., Mochizuki, Y., Komeiji, Y., Okiyama, Y., and Fukuzawa, K. (2014). Electron-correlated fragment-molecular-orbital calculations for biomolecular and nano systems. Physical Chemistry Chemical Physics 16, 10310–10344.

Taylor, S.S., Knighton, D.R., Zheng, J., Ten Eyck, L.F., and Sowadski, J.M. (1992). Structural framework for the protein kinase family. Annu Rev Cell Biol 8, 429–462.

Traut, T. (1994). Physiological concentrations of purines and pyrimidines. Molecular and Cellular Biochemistry 140, 1–22.

Vadas, O., Burke, J.E., Zhang, X., Berndt, A., and Williams, R.L. (2011). Structural Basis for Activation and Inhibition of Class I Phosphoinositide 3-Kinases. Science Signaling 4, re2.

Wang, Z., and Cole, P.A. (2014). Catalytic mechanisms and regulation of protein kinases. Methods Enzymol 548, 1–21.

Wennerberg, K., Rossman, K.L., and Der, C.J. (2005). The Ras superfamily at a glance. Journal of Cell Science 118, 843.

