## Supplemental materials for "Activity-specificity trade-off gives PI5P4Kβ a nucleotide preference to function as a GTP-sensing kinase"

### Supplemental lists

List of PDB identification numbers used in the structural analysis of adenine and guanine nucleotide complexes of kinases and G-proteins, shown in Figure 2.

***List of kinases in complex with adenine nucleotides***

1AD5, 1ATP, 1B38, 1CDK, 1CJA, 1CM8, 1CSN, 1DAW, 1DS5, 1E8X 1FIN, 1FQ1, 1GOL, 1GY3, 1H1W, 1HCK, 1I44, 1IA9, 1IAH, 1IG1 1IR3, 1J1B, 1J1C, 1JBP, 1JKK, 1JKL, 1JNK, 1JPA, 1JQH, 1JST, 1JWH, 1K3A, 1L3R, 1LP4, 1MP8, 1MQ4, 1MQB, 1MRU, 1NY3, 1O6K, 1O6L, 1O6Y, 1OL5, 1OL6, 1OL7, 1PHK, 1PJK, 1PKG, 1PYX, 1Q24, 1Q8Y, 1Q97, 1Q99, 1QL6, 1QMZ, 1QPC, 1RDQ, 1S9I, 1S9J, 1TQM, 1TQP, 1TZD, 1U54, 1U5R, 1UA2, 1W2C, 1W2D, 1WBP, 1XR1, 1YXT, 1YXU, 1Z2N, 1Z2O, 1Z2P, 1ZAO, 1ZAR, 1ZP9, 1ZTH, 1ZY5, 1ZYD, 2A19, 2ACX, 2AQX, 2B9F, 2B9H, 2B9I, 2B9J, 2BIY, 2BZK, 2C6D, 2CCH, 2CCI, 2CJM, 2DWB, 2EB3, 2F9G, 2G2F, 2G2I, 2GS7, 2HEN, 2IJM, 2ITN, 2ITV, 2ITX, 2IVT, 2J0L, 2OID, 2OZO, 2P0C, 2P55, 2PHK, 2PMI, 2PML, 2PVF, 2PVR, 2PVY, 2PWL, 2PY3, 2PYW, 2PZ5, 2PZP, 2PZR, 2Q0B, 2Q7D, 2QB5, 2QCS, 2QO7, 2QO9, 2QOC, 2QOQ, 2QUR, 2RIO, 2SRC, 2V55, 2V8Q, 2V92, 2V9J, 2VED, 2VWI, 2W4J, 2W4K, 2W5A, 2W5B, 2WQE, 2WQN, 2WTK, 2XAL, 2XAM, 2XAN, 2XRW, 2XS0, 2XUU, 2Y8L, 2Y8Q, 2Y9Q, 2YA3, 2YAA, 2YAB, 2Z7Q, 2ZV8, 3A7H, 3A7J, 3A8W, 3A99, 3AKK, 3AKL, 3ALN, 3ALO, 3BEG, 3BFV, 3BLQ, 3BRB, 3BU5, 3C0H, 3C4W, 3C4X, 3C4Z, 3C50, 3C51, 3CLY, 3COK, 3D5W, 3DAK, 3DFC, 3DKC, 3DLS, 3DLZ, 3DQW, 3DQX, 3DV3, 3DY7, 3E8N, 3EH9, 3EHA, 3EQB, 3EQC, 3EQD, 3EQG, 3EQH, 3EQI, 3F5G, 3F5U, 3F61, 3FHI, 3FJQ, 3FXX, 3FZP, 3G2F, 3G51, 3GC0, 3GNI, 3GQI, 3GT8, 3GU4, 3GU5, 3GU6, 3GU7, 3HKO, 3HMN, 3HRC, 3HRF, 3HX4, 3IDB, 3IDC, 3IGO, 3IS5, 3JBZ, 3JUH, 3KEX, 3KMW, 3KN5, 3KU2, 3LCT, 3LIJ, 3LJ0, 3LKM, 3LLA, 3LLT, 3LMG, 3LMH, 3LMI, 3MBL, 3MFR, 3MFS, 3MFU, 3MIA, 3NIE, 3NIZ, 3NSZ, 3NYO, 3O7L, 3O8L, 3O8N, 3ORI, 3ORK, 3ORL, 3ORM, 3ORN, 3ORO, 3ORP, 3ORT, 3OS3, 3P23, 3PDT, 3PFQ, 3PP1, 3PVB, 3Q4Z, 3Q53, 3Q5I, 3QAL, 3QAM, 3QBW, 3QC9, 3QHR, 3QHW, 3REP, 3SLS, 3T4N, 3T54, 3T7A, 3T8O, 3T99, 3T9A, 3T9B, 3T9C, 3T9D, 3T9E, 3T9F, 3TDH, 3TEI, 3TNQ, 3TPT, 3U87, 3UDS, 3UDT, 3UDZ, 3UIM, 3V01, 3V04, 3VJN, 3VJO, 3VVH, 3W8Q, 3WIG, 3WOW, 3X01, 3X03, 3X05, 3X07, 3X09, 3X0B, 3X2U, 3X2V, 3X2W, 3ZXT, 4A06, 4A07, 4AN2, 4AN3, 4AN9, 4ANB, 4AQK, 4ARK, 4AW0, 4AW1, 4AXD, 4AXE, 4AXF, 4BFM,

4BL1, 4BN1, 4BTJ, 4C2W, 4C3P, 4C3R, 4CEG, 4CFE, 4CFF, 4CFH, 4CRS, 4CT1, 4CT2, 4CXA, 4DEE, 4DFX, 4DG0, 4DG3, 4DH1, 4DH3, 4DH5, 4DH7, 4DH8, 4DIN, 4DN5, 4EAG, 4EAI, 4EAJ, 4EAK, 4EAL, 4EKK, 4EOJ, 4EOM, 4EOO, 4EOQ, 4EQM, 4F0F, 4F1M, 4F1O, 4F99, 4F9A, 4FG7, 4FG8, 4FG9, 4FI1, 4FIE, 4FIF, 4FIG, 4FL1, 4FL2, 4FL3, 4FMQ, 4FVQ, 4FVR, 4GT3, 4GVA, 4GVJ, 4GYI, 4H3P, 4H3Q, 4HN2, 4HND, 4HNE, 4HPT, 4HPU, 4I3Z, 4I94, 4IAC, 4IAD, 4IAF, 4IAI, 4IAK, 4IAY, 4IAZ, 4IB0, 4IB1, 4IB3, 4IC7, 4IEB, 4IFC, 4II5, 4IIR, 4IMY, 4IX4, 4IX5, 4IX6, 4IZ5, 4J95, 4J96, 4J97, 4J98, 4J99, 4JDI, 4JQE, 4JRN, 4JSP, 4JSV, 4K2R, 4K33, 4KKE, 4KQB, 4KRC, 4LGD, 4LMN, 4LQP, 4LQQ, 4LQS, 4LRJ, 4LV7, 4M15, 4MNE, 4MO4, 4MO5, 4MWH, 4NH1, 4NIF, 4NM0, 4NM3, 4NM5, 4NM7, 4NST, 4NTT, 4NU1, 4NZM, 4NZN, 4NZO, 4O21, 4O27, 4O4D, 4O4E, 4O4F, 4OH4, 4OTP, 4PDS, 4PL3, 4PL4, 4PL5, 4PLA, 4Q4C, 4Q4D, 4Q5H, 4Q5J, 4QFG, 4QFR, 4QFS, 4QML, 4QNY, 4QPM, 4QTD, 4R8Q, 4RER, 4REW, 4RIW, 4RIX, 4RIY, 4RQK, 4RQV, 4RRV, 4S32, 4S33, 4S34, 4TND, 4U40, 4U7Z, 4U80, 4U81, 4UAK, 4UEU, 4UX9, 4UYA, 4WB5, 4WB6, 4WB7, 4WB8, 4WBB, 4WTV, 4WW5, 4WW7, 4WW9, 4X6R, 4XBR, 4XF6, 4XF7, 4XH0, 4XHG, 4XLV, 4XW4, 4XW5, 4XW6, 4XX9, 4Y0X, 4Y12, 4Y5Q, 4YC4, 4YSJ, 4YZD, 4Z9L, 4ZMF, 4ZS4, 4ZSE, 5ACK, 5AR3, 5AWM, 5BYA, 5BYB, 5C03, 5CE3, 5CKW, 5CNN, 5CNO, 5CSH, 5CU6, 5CVF, 5CVH, 5D41, 5D9H, 5DBX, 5DGH, 5DGI, 5DMZ, 5DN3, 5DNR, 5DOS, 5DR2, 5DRD, 5DT3, 5DT4, 5DYJ, 5DZC, 5E3T, 5E3U, 5E92, 5E9E, 5EFQ, 5EG3, 5FG8, 5G15, 5G1X, 5HNV, 5HU3, 5HVJ, 5HVK, 5I0N, 5I35, 5I4N, 5I9V, 5I9W, 5ISO, 5JR7, 5KHW, 5KQ5, 5L6W, 5L8J, 5L8K, 5L8L, 5LI1, 5LI9, 5LIH, 5LPB, 5LPV, 5LPY, 5LPZ, 5LVO, 5LVP, 5LXM, 5M06, 5MOE, 5MP8, 5MPJ, 5MRD, 5MW8, 5MWL, 5NCL, 5NG0, 5NZZ, 5O0Y, 5O26, 5OBJ, 5ODT, 5OOP, 5OQU, 5ORJ, 5ORK, 5ORL, 5ORN, 5ORO, 5ORP, 5ORR, 5ORS, 5ORT, 5ORV, 5ORW, 5ORX, 5ORY, 5ORZ, 5OS0, 5OS1, 5OS2, 5OS3, 5OS4, 5OS5, 5OS6, 5OS7, 5OS8, 5OSD, 5OSE, 5OSF, 5OSL, 5OSP, 5OSR, 5OTR, 5T5T, 5TF9, 5UFU, 5UGL, 5UGX, 5UHN, 5UI0, 5UPK, 5V60, 5W2H, 5W2I, 5W7T, 5WDY, 5WE8, 5WNI, 5WNO, 5X17, 5X1Q, 5X1S, 5X1T, 5X2A, 5XD6, 5XVU, 5XZV, 5YT3, 6AC9, 6AO5, 6B1U, 6B2E, 6BCU, 6BCX, 6BG2, 6BXI, 6C7Y, 6C83, 6C9F, 6C9G, 6C9H, 6C9J, 6CMW, 6CN2, 6CPF, 6CQD, 6E4T, 6E4U, 6E4W, 6EGF, 6EH2, 6EM6, 6EM7, 6EMA, 6EQ9, 6ESA, 6F3F, 6FD3, 6FJK, 6FL3, 6FL8, 6GFG, 6GFH, 6GMD

***List of G-proteins in complex with guanine nucleotides***

1CC0, 1D5C, 1F6B, 1FZQ, 1KAO, 1KY3, 1M7B, 1MKY, 1N6K, 1OIX, 1R8S, 1SVI, 1T91, 1TAD, 1U0L, 1U8Z, 1VG1, 1WMS, 1YRB, 1Z0A, 1Z0F, 1Z0I, 1Z0J, 1Z2A, 1ZBD, 1ZCA, 1ZCB, 2A5D, 2A5J, 2ATV, 2BCG, 2BMD, 2C03, 2CE2, 2CLS, 2CXX, 2DPX, 2E9S, 2EFH, 2ERX, 2ERY, 2F7S, 2F9L, 2FN4, 2FQX, 2FV8, 2G3Y, 2G77, 2GF0, 2GF9, 2HT6, 2HUP, 2IL1, 2IYL, 2J0V, 2J1L, 2KSQ, 2NZJ, 2O52, 2ODE, 2Q3H, 2QTH, 2RHD, 2WIB, 2WJG, 2WJH, 2WKQ, 2YV5, 2YWH, 3A6P, 3BH7, 3BWD, 3C5C, 3CBQ, 3CLV, 3CNO, 3DZ8, 3EC1, 3GJ0,

3IEU, 3KKQ, 3LVR, 3M1I, 3O47, 3P27, 3P32, 3REF, 3RWO, 3SFV, 3SYN, 3T1O, 3W6P, 3X1W, 4ARZ, 4DCU, 4DJT, 4FID, 4KLZ, 4KU4, 4LC1, 4LHV, 4M8N, 4MIT, 4QXA, 4WNR, 4Z8Y, 5C2K, 5C4M, 5DN8, 5G53, 5HCI, 5JCP, 5M04, 5O33, 5UB8, 5UF8, 5VCU, 5WDS, 5X4B, 5XC5, 5YMX, 6AU6, 6BSX, 6D4G, 6DI7, 6EKK, 6FF8, 6G0Z

### Supplemental Figures

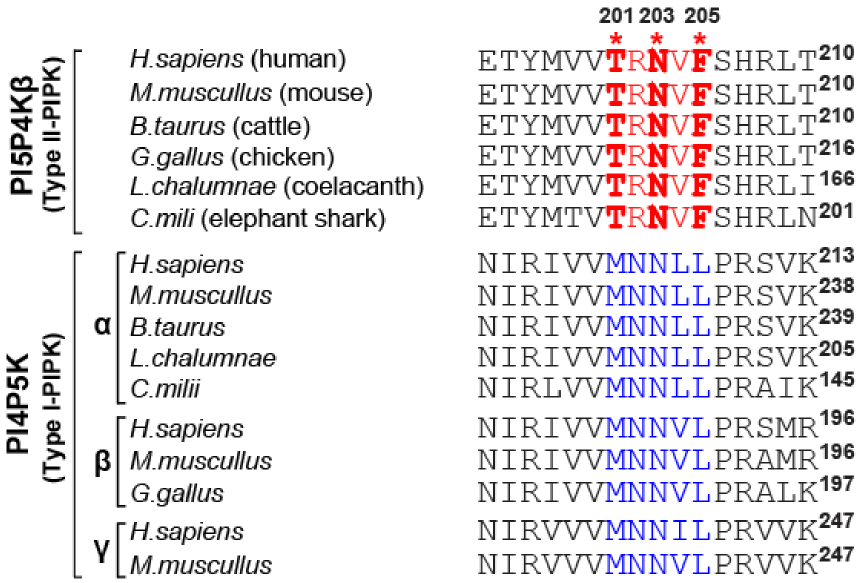

**Fig. S1 Sequence alignment of PI5P4K and PI4P5K family proteins.** The GTP-recognizing TRNVF sequences and the ATP-recognizing MNNψL sequences are colored in red and blue, respectively. The positions Thr-201, Asn-203, and Phe-205 in human PI5P4Kβ are indicated by asterisks (*).

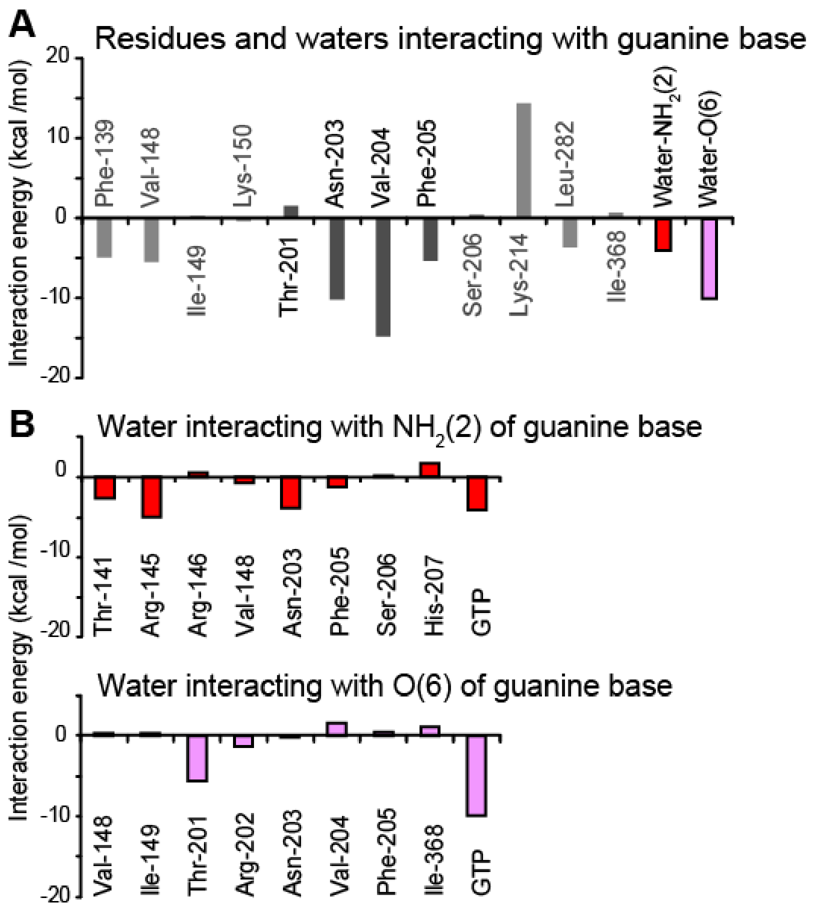

**Fig. S2 Fragment molecular orbital (FMO) calculation of the PI5P4Kβ-GTP complex**

(A) The energetic contributions of each residue to PI5P4Kβ-guanine base interaction are indicated. The energetic contributions of two water molecules that are bound to the $NH_2(2)$ and O(6) positions of the guanine base moieties are also shown. (B) The energetic contributions of each residue of PI5P4Kβ and GTP to the interaction with water molecules that are bound to the (top) $NH_2(2)$ and (bottom) O(6) positions, respectively, of guanine base moieties are indicated.

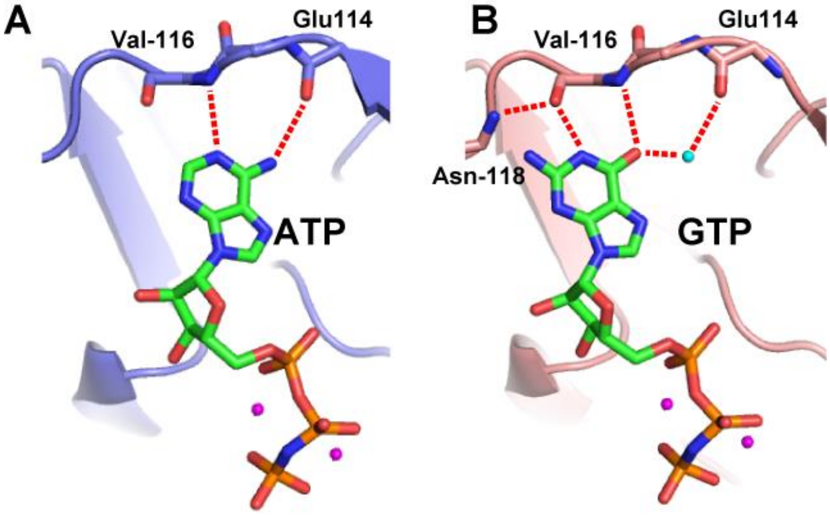

**Fig. S3 Interaction of (A) ATP and (B) GTP with CKII.** Red dotted lines represent the hydrogen bonds between the triphosphorylated nucleotides and CKII. The figure was generated from the structure of *Zea mays* CKIIα under the accession codes 1DAW (CKIIα-AMPPNP complex) and 1DAY (CKIIα- GMPPNP complex).

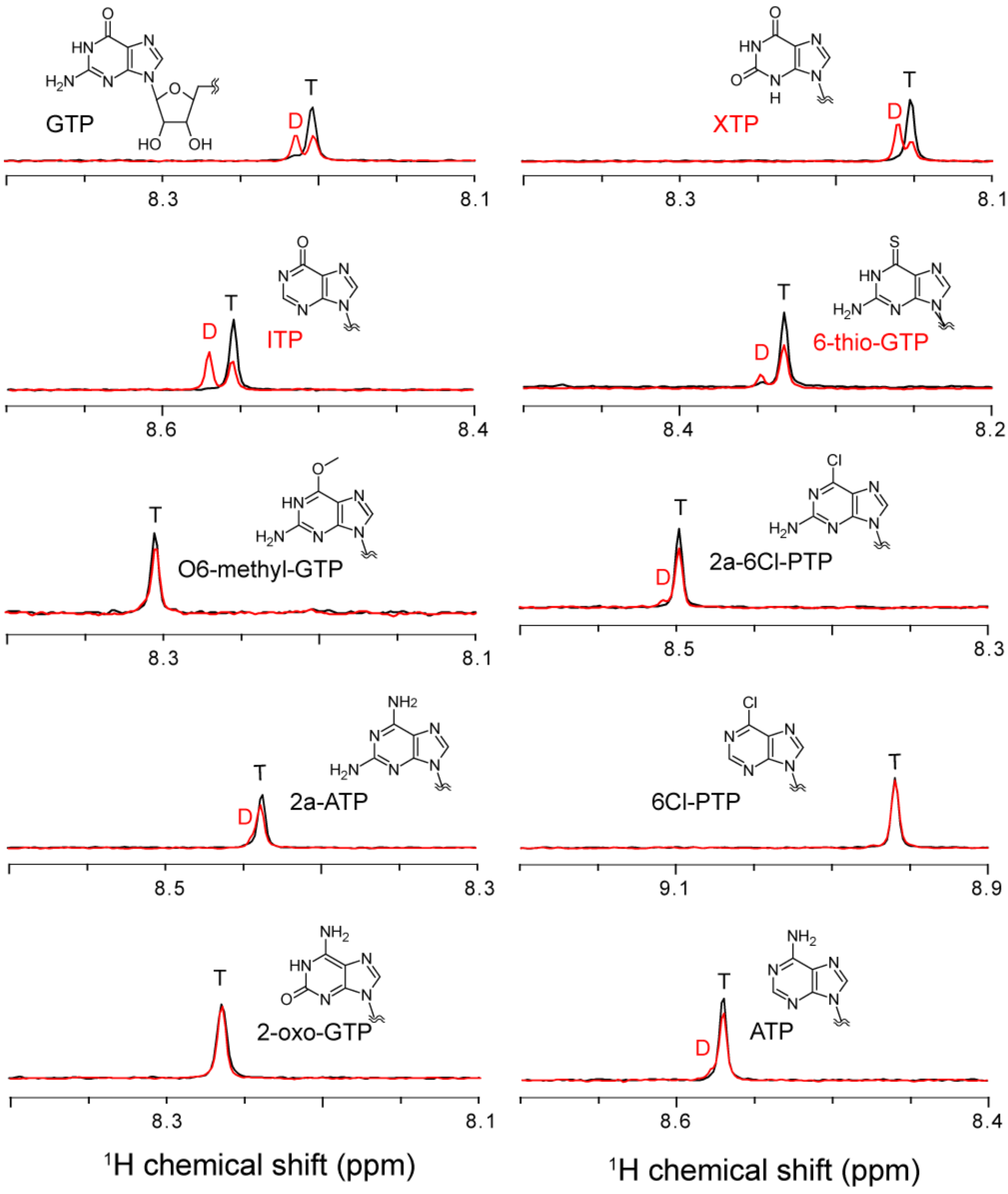

**Fig. S4 NMR-based PNT hydrolysis assay**. The recombinant PI5P4Kβ was incubated with 250 μM PNTs. The protein concentration was fixed to 2 μM. The intensities of H8 protons of dephosphorylated (D)/triphosphorylated (T) nucleotide signals are quantified from the spectra.

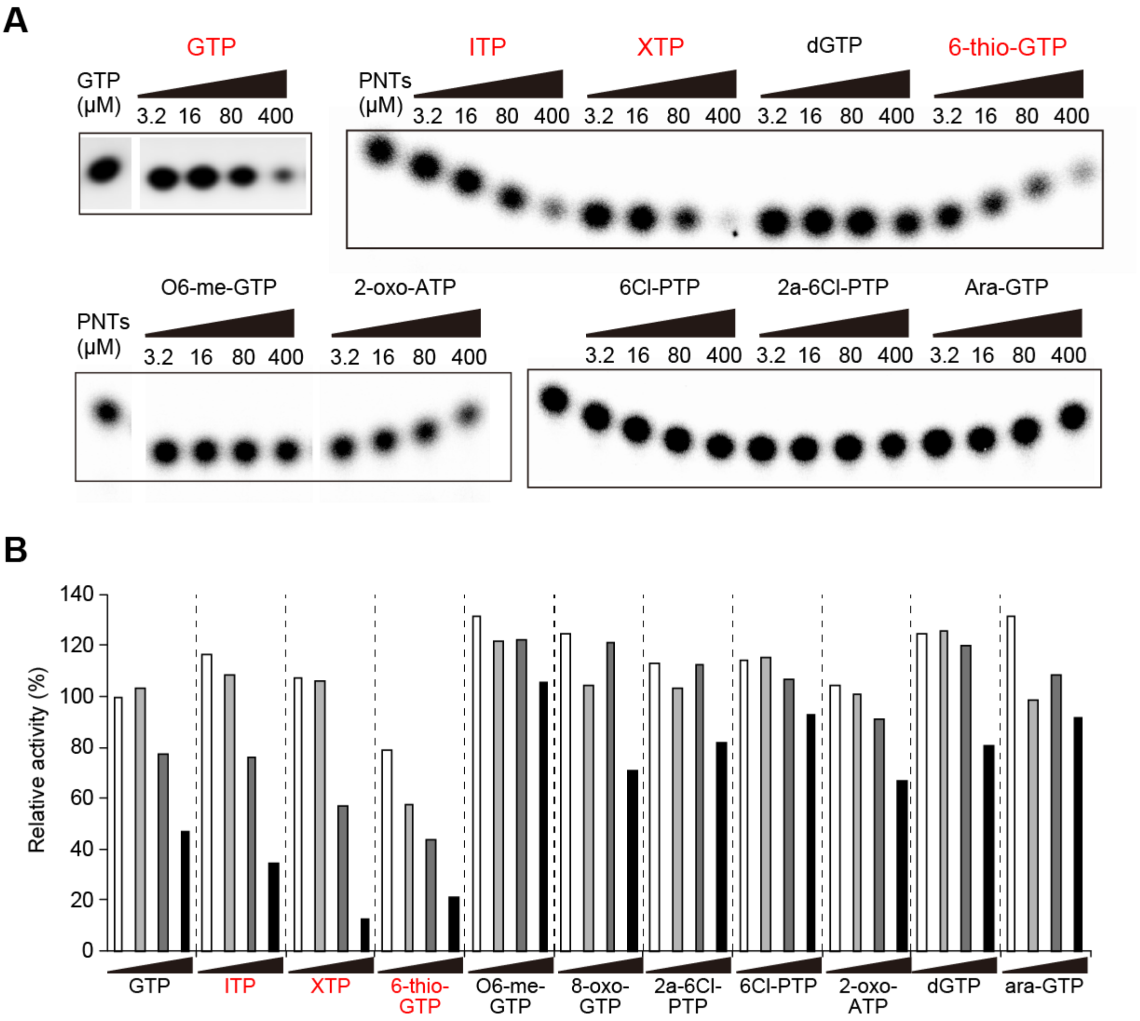

**Fig. S5 Inhibition of the $^{32}$P-GTP-dependent kinase activity of PI5P4Kβ by cold PNTs.** (A) The *in vitro* kinase assay was performed as previously described (Davis et al., 2013; Rameh et al., 1997). The kinase reaction was carried out in a total of 50 μl of reaction buffer (50 mM HEPES (pH 7.4), 0.2 mM EGTA, and 10 mM $MgCl_2$) containing 20 μM of PI(5)P (d-myo-phosphatidylinositol 5-phosphate diC16) that was suspended by sonication. 250 μM of γ-$^{32}$P radiolabeled GTP (Perkin Elmer) was incubated with 1 μg of recombinant PI5P4Kβ for 10 min at room temperature. Unless otherwise indicated, 20 μM of PI(5)P was used with 3 μM 1,2-dipalmitoyl-phosphatidylserine as the basal lipid. Phosphoinositides were extracted by a methanol/chloroform (1/1, v/v) mix and subjected to a thin-layer chromatography (TLC) assay using heat-activated 2% oxaloacetate-coated silica gel 60 plates (EMD Chemicals Inc., Billerica,

MA, USA). As for the solvent, 1-propanol/2 M acetic acid (65/35, v/v) was used. The radiolabeled $PI(4,5)P_2$ was quantified with a phosphorimager (Typhoon Trio; GE Healthcare). The data were analyzed and graphed by Prism 6 (GraphPad Software) or KaleidaGraph 4.1 (Synergy Software). In competition assay, 250 μM of $^{32}$P-labeled GTP was mixed with 3.2 - 400 μM triphosphorylated nucleotides, and the PI(5)P phosphorylation by GTP was monitored by quantifying the amount of radiolabeled $PI(4,5)P_2$. (B) Relative kinase activities in the presence of each PNT against those without them (*i.e.,* only $^{32}$P-GTP condition) are shown. The concentrations of PNTs are 3.2, 16, 80, and 400 μM (see panel A). PNTs that decreased the $^{32}$P GTP-dependent activity to less than 50% of the control value are indicated in red.

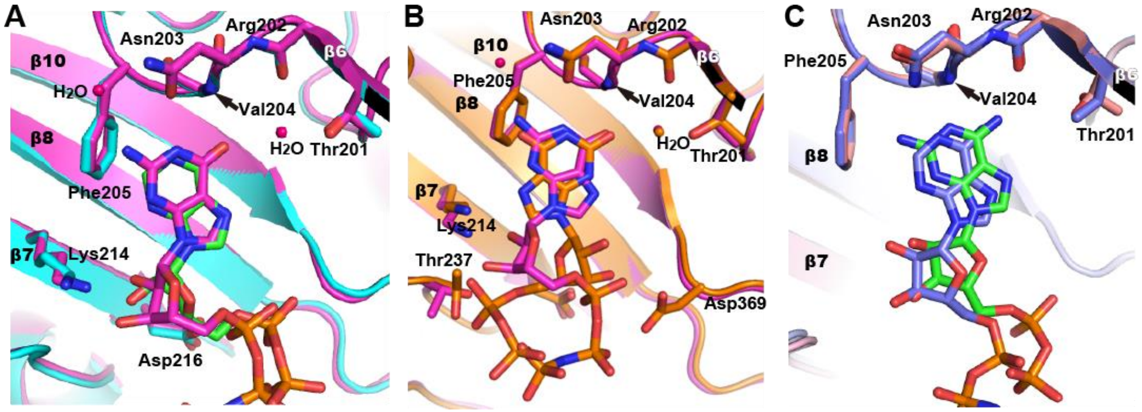

**Fig. S6 Overlay of the ITP, XTP, and 2a-ATP binding modes with the GTP or ATP binding modes.** (A) the PI5P4Kβ-ITP complex and (B) the PI5P4Kβ-XTP complexes are overlaid with the PI5P4Kβ-GTP complex (GTP: magenta). For (C) the PI5P4Kβ-2a-ATP complex, the PI5P4Kβ-ATP complex (ATP: purple) was overlaid.

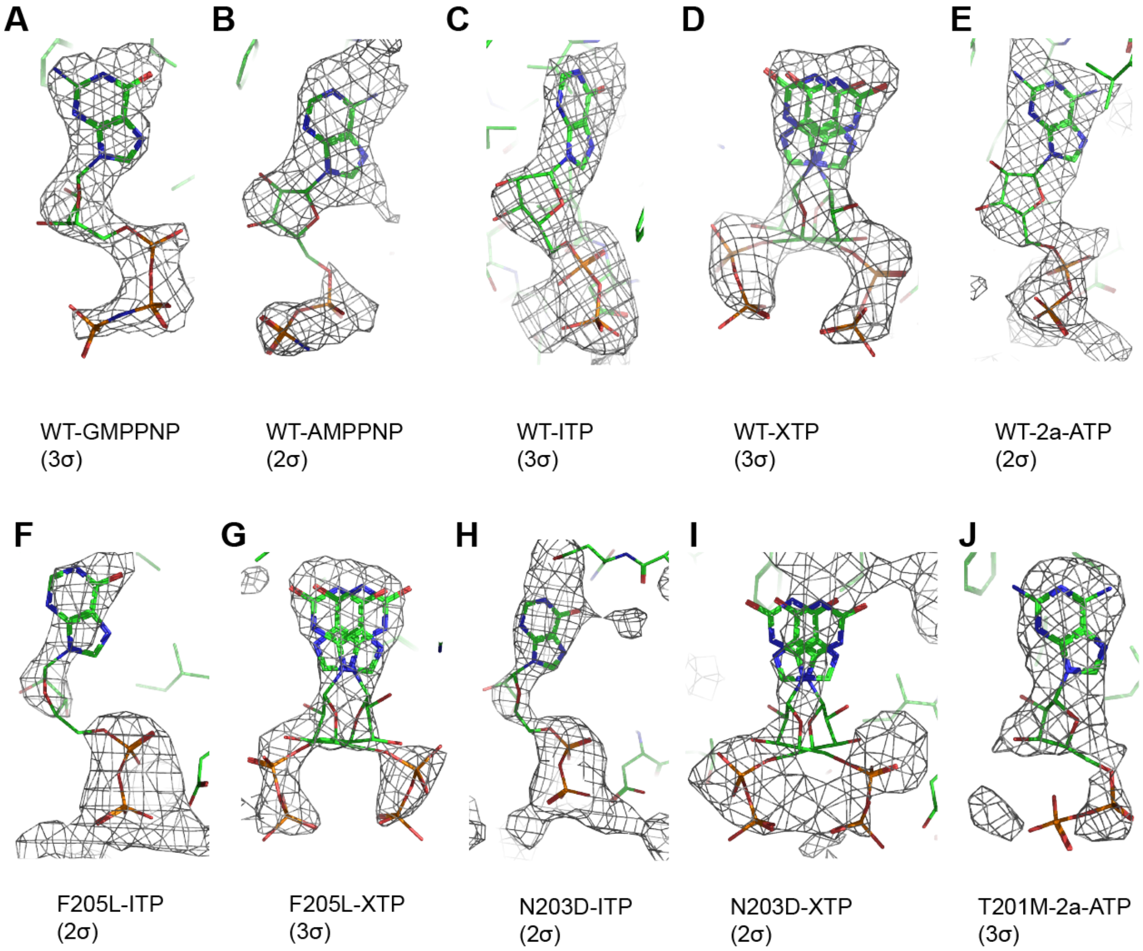

**Fig. S7 Simulated annealing $mF_o$-$DF_c$ omit maps for bound nucleotides in PI5P4Kβ.** The positive $mF_o$-$DF_c$ electron densities of the indicated nucleotides in the active site of PI5P4Kβ were contoured at the indicated levels (2-3 σ) and shown in the same direction. These densities show that the nucleotides and their positions were unambiguously identified in the PI5P4Kβ-nucleotide complexes.

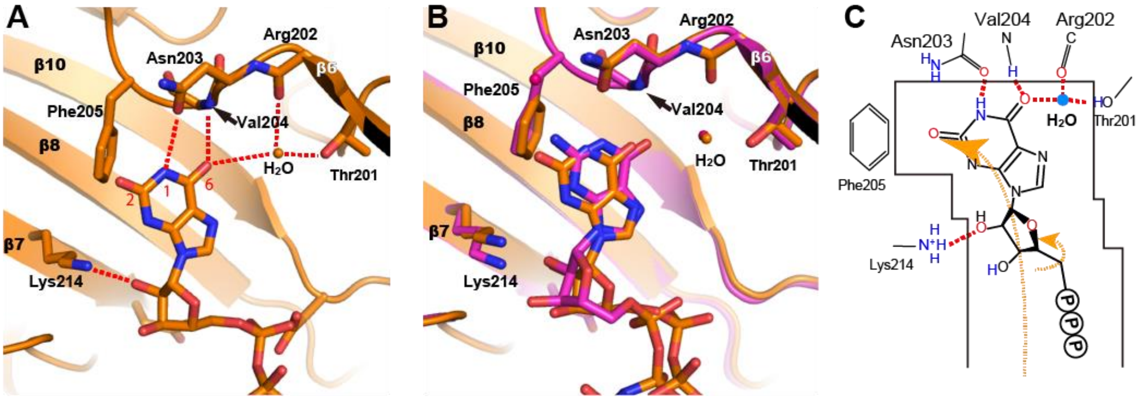

**Fig. S8 Interaction of XTP with PI5P4Kβ in GTP-binding mode.** (A) Binding of XTP by PI5P4Kβ in the GTP-binding mode. The red dotted lines represent the hydrogen bonds between the triphosphorylated nucleotides and PI5P4Kβ. (B) The PI5P4Kβ-XTP complex in GTP-binding mode was overlaid with the PI5P4Kβ-GTP complex. GTP is shown in magenta. (C) Schematic representations of the GTP-binding mode of XTP.

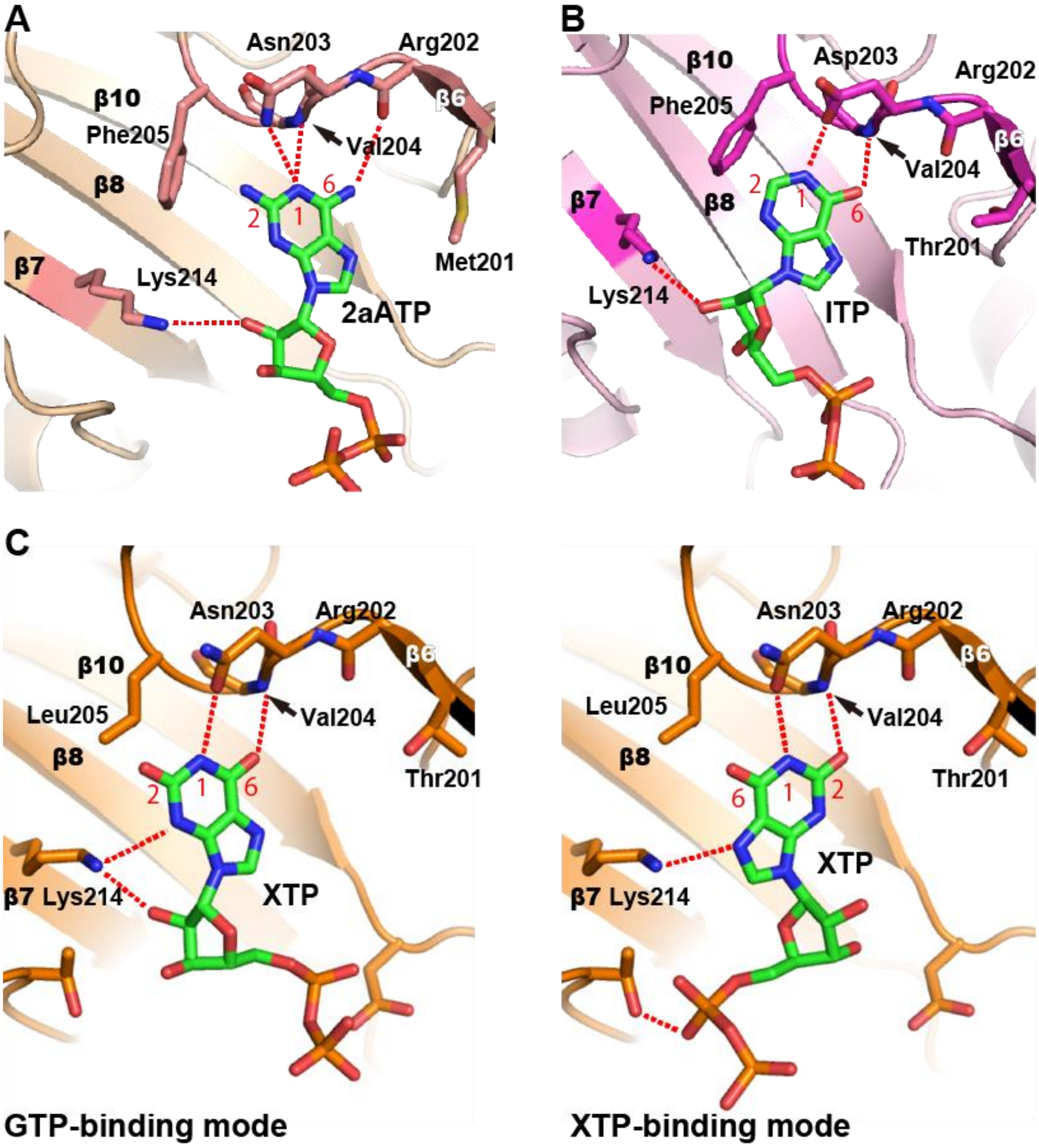

**Fig. S9 Interaction of PNTs with PI5P4Kβ mutants.** (A) 2a-ATP binding to PI5P4Kβ$^{T201M}$ in ATP-binding mode. (B) ITP binding to PI5P4Kβ$^{N203D}$ in GTP-binding modes. (C) XTP binding to PI5P4Kβ$^{F205L}$ in (left) GTP- and (right) XTP-binding modes. The red dotted lines represent the hydrogen bonds between the triphosphorylated nucleotides and PI5P4Kβ.

**Table S1 Data collection and refinement statistics**

| | WT-GMPPNP | WT-AMPPNP | WT-ITP | WT-XTP |
|---|---|---|---|---|
| **Data collection** | | | | |
| Space group | $C222_1$ | $C222_1$ | $C222_1$ | $C222_1$ |
| Cell dimensions | | | | |
| *a*, *b*, *c* (Å) | 109.8 | 108.6 | 109.2 | 109.9 |
| | 184.6 | 183.0 | 186.9 | 185.2 |
| | 105.9 | 106.0 | 104.7 | 105.8 |
| Resolution (Å) | 94.36-2.70 | 91.49-2.55 | 69.69-2.65 | 70.47-2.60 |
| | (2.85-2.70) | (2.69-2.55) | (2.79-2.65) | (2.74-2.60) |
| $R_{merge}$ (%) | 4.5 (90.8) | 3.0 (82.1) | 4.7 (84.5) | 3.4 (92.9) |
| $I/\sigma I$ | 35.3 (2.7) | 43.6 (2.5) | 23.6 (3.1) | 40.0 (2.9) |
| Completeness (%) | 99.3 (99.9) | 98.3 (99.1) | 99.9 (99.9) | 99.9 (99.9) |
| Redundancy | 7.2 (6.9) | 7.2 (6.3) | 8.6 (9.1) | 8.7 (9.0) |
| **Refinement** | | | | |
| Resolution (Å) | 37.69-2.70 | 53.18-2.55 | 45.65-2.65 | 38.10-2.60 |
| No. reflections | 29713 | 34181 | 31448 | 33522 |
| $R_{work}/R_{free}$ (%) | 20.8/27.7 | 22.8/28.2 | 22.8/26.6 | 22.0/25.8 |
| No. atoms | | | | |
| Protein | 4742 | 4708 | 4673 | 4656 |
| Ligand/ion | 96 | 76 | 85 | 144 |
| Water | 5 | 5 | 0 | 7 |
| B-factors | | | | |
| Protein | 88.3 | 98.4 | 89.9 | 89.9 |
| Ligand/ion | 123.6 | 138.3 | 96.5 | 119.7 |
| Water | 64.1 | 80.7 | - | 74.1 |
| R.m.s deviations | | | | |
| Bond lengths (Å) | 0.009 | 0.008 | 0.009 | 0.009 |
| Bond angles (º) | 0.995 | 1.021 | 1.018 | 1.200 |
| PDB ID | 6K4G | 6K4H | 7C54 | 7C55 |

**Table S1 Data collection and refinement statistics (continued)**

| | WT-2a-ATP | F205L-ITP | F205L-XTP | N203D-ITP |
|---|---|---|---|---|
| **Data collection** | | | | |
| Space group | $C222_1$ | $C222_1$ | $C222_1$ | $C222_1$ |
| Cell dimensions | | | | |
| *a*, *b*, *c* (Å) | 109.8 | 109.3 | 109.6 | 109.1 |
| | 185.9 | 185.5 | 186.2 | 186.4 |
| | 106.2 | 105.7 | 105.7 | 106.3 |
| Resolution (Å) | 48.78-3.10 | 48.55-2.80 | 48.63-2.80 | 48.53-2.95 |
| | (3.27-3.10) | (2.95-2.80) | (2.95-2.80) | (3.11-2.95) |
| $R_{merge}$ (%) | 6.4 (89.6) | 6.2 (89.0) | 6.1 (99.6) | 6.3 (95.6) |
| $I/\sigma I$ | 24.6 (3.0) | 28.9 (2.7) | 25.5 (2.5) | 26.8 (2.7) |
| Completeness (%) | 99.9 (100.0) | 99.9 (99.7) | 99.7 (98.4) | 99.9 (100.0) |
| Redundancy | 8.8 (9.2) | 8.9 (8.3) | 8.9 (8.5) | 9.0 (9.4) |
| **Refinement** | | | | |
| Resolution (Å) | 46.12-3.10 | 45.94-2.80 | 47.21-2.80 | 35.69-2.95 |
| No. reflections | 20085 | 26819 | 26898 | 23179 |
| $R_{work/}$ $R_{free}$ (%) | 21.3/25.6 | 21.6/27.8 | 21.7/26.1 | 21.4/26.8 |
| No. atoms | | | | |
| Protein | 4574 | 4677 | 4669 | 4633 |
| Ligand/ion | 60 | 84 | 116 | 85 |
| Water | 0 | 1 | 2 | 0 |
| B-factors | | | | |
| Protein | 109.3 | 85.3 | 87.7 | 97.1 |
| Ligand/ion | 147.1 | 111.4 | 139.8 | 107.1 |
| Water | - | 62.8 | 62.3 | - |
| R.m.s deviations | | | | |
| Bond lengths (Å) | 0.010 | 0.009 | 0.009 | 0.009 |
| Bond angles (º) | 1.227 | 1.096 | 1.106 | 1.106 |
| PDB ID | 7C56 | 7C57 | 7C58 | 7C59 |

**Table S1 Data collection and refinement statistics (continued)**

| | N203D-XTP | T201M-2a-ATP |
|---|---|---|
| **Data collection** | | |
| Space group | $C222_1$ | $C222_1$ |
| Cell dimensions | | |
| *a*, *b*, *c* (Å) | 107.5 | 109.5 |
| | 185.6 | 184.9 |
| | 107.0 | 106.7 |
| Resolution (Å) | 48.04-3.55 | 47.97-3.05 |
| | (3.74-3.55) | (3.21-3.05) |
| $R_{merge}$ (%) | 6.1 (87.4) | 7.0 (90.9) |
| $I/\sigma I$ | 23.0 (2.4) | 24.2 (2.9) |
| Completeness (%) | 99.8 (99.9) | 99.9 (100.0) |
| Redundancy | 8.8 (8.4) | 8.8 (8.6) |
| **Refinement** | | |
| Resolution (Å) | 33.29-3.55 | 46.22-3.05 |
| No. reflections | 13257 | 20989 |
| $R_{work/}$ $R_{free}$ (%) | 24.9/30.0 | 20.9/27.5 |
| No. atoms | | |
| Protein | 4317 | 4605 |
| Ligand/ion | 88 | 60 |
| Water | 0 | 0 |
| B-factors | | |
| Protein | 165.4 | 101.0 |
| Ligand/ion | 179.5 | 118.2 |
| Water | - | - |
| R.m.s deviations | | |
| Bond lengths (Å) | 0.003 | 0.010 |
| Bond angles (º) | 0.719 | 1.196 |
| PDB ID | 7C5A | 7C5B |

Each structure was determined from single-crystal diffraction.
The highest resolution shell is shown in parentheses.
